# Circulating extracellular vesicles can transport stress signals to the male germline

**DOI:** 10.1101/2025.03.21.644551

**Authors:** Anar Alshanbayeva, Leonard C. Steg, Alaa Othman, Francesca Manuella, Rodrigo G. Arzate-Mejía, Nicola Zamboni, Isabelle M. Mansuy

## Abstract

Extracellular vesicles (EVs) play a key role in cell-cell communication by transporting bioactive molecules from donor to recipient cells across the body. While their involvement in somatic cells communication is well described, somatic-to-germ cells communication remains understudied. We show that small EVs (sEVs) can vehicle signals of early life stress (ELS) from circulation to sperm consequentially for the offspring. In mice, ELS persistently modifies RNA, lipids and metabolites in plasma sEVs. Chronic injection of plasma sEVs from ELS-exposed males alters the sperm transcriptome and is reflected in the transcriptome of embryos. This leads to metabolic dysfunctions in the adult offspring. These findings highlight a role of circulating sEVs in soma-to-germline communication relevant for the intergenerational transmission of ELS effects.

## Introduction

Intercellular communication is a fundamental biological process necessary for development and cellular functions in multicellular organisms. It allows cells to exchange information within local microenvironments and over long distances. Classical modes of communication of cells in close proximity involve exocytosis and/or sharing of small cytoplasmic components through gap junctions. For cells that are far apart, endocrine signaling allows communication through specific factors secreted in the bloodstream (*1, 2*). Further to these well-known processes, a recently identified mode of cell-to-cell communication involves extracellular vesicles (EVs). EVs are small cargo-bearing vesicles, released by nearly all cells in the extracellular space and in circulation, that enable the transfer of biological material over short and long distances (*3, 4*). EVs have different sizes and modes of secretion and include small EVs (sEVs) (40-160nm) released by exocytosis, microvesicles (150nm-1um) formed by budding, and apoptotic bodies (0,1-5um) formed by fragmentation during apoptosis (*5*). They carry functional molecules such as nucleic acids, proteins, lipids and metabolites and play important roles in physiological, immunological and metabolic functions (*6, 7*). An important property of EVs is their intrinsic ability to cross cellular barriers (*8*). Thus, EVs released from a given tissue can be vehicled and transfer their cargo to another tissue, thereby inducing cellular changes in distant cells. For instance, EVs from adipose tissue can reach liver and regulate gene expression in hepatocytes (*9*), while EVs from peripheral blood can cross the blood-brain barrier and modulate neuroinflammation in the brain (*10*). In pathological states such as cancer, EVs released from lung, liver or breast tumor cells can modulate immune cells and lymphoid components in the tumor microenvironment (*11*). EVs have also been implicated in mental disorders, although little is known (*12*).

While the role of EVs in somatic cells communication has been well studied (*13*), whether circulating EVs can reach gonads and provide signals to germ cells remains unexplored. This is a critical question because germ cells can be modified by peripheral factors (*14*) and environmental exposure, in some cases with consequences for the offspring (*15, 16*). Rodent studies have shown that maturing sperm cells can acquire bioactive components from EVs in epididymis (*17*) and that the RNA content of epididymosomes can be altered by paternal stress or poor diet and influence sperm RNA cargo (*18, 19*).

We examined if circulating EVs can be modified by early life stress (ELS), a condition with widespread and durable effects on health in humans and animals and that can affect reproductive cells and the offspring (*20*–*23*). Using a mouse model of chronic stress in postnatal life with transgenerational phenotypes (*24, 25*), we show that ELS alters the cargo and composition of sEVs in plasma in adulthood. We tested if plasma sEVs can influence sperm by intravenously injecting plasma sEVs from males exposed to ELS into naïve males. We observed that plasma sEVs modify the sperm transcriptome of injected males, which correlates with altered transcriptional programs and metabolic dysregulation in the offspring. These results suggest that circulating sEVs carry physiological signals of stress from soma to the germline and are likely involved in father-to-offspring transfer of paternal experiences.

## Results

### Circulating sEVs have a rich RNA, lipid and metabolite content

We isolated circulating EVs by fractionating blood plasma from adult male mice using size-exclusion chromatography (SEC), a method that separates biological components including EVs by size (*26*) (**fig. S1A**). We obtained 13 fractions (1-13) and in two of them (4-5), we identified particles with typical markers (*27*), size (50-120nm in diameter) and morphology of sEVs and no ApoE1 contaminant (**Fig. 1A-C and fig. S1B)**. In these 2 fractions, RNA sequencing revealed complex RNA populations with a majority of reads mapping to transfer RNAs (tRNAs), microRNAs (miRNAs) and protein coding regions, as previously reported (*9, 27*) (**Fig. 1D**). Several miRNAs such as miR-30a and miR-10b (*27*) – known to be enriched in sEVs – were abundant, while miRNAs carried by HDL particles (*28*) were scarce, confirming the specificity of our sEVs preparations (**fig. S2A**). Analyses of metabolites showed that carbohydrates, fatty acyls, benzoids and steroids among other metabolite sets are enriched in sEVs preparations (**Fig. 1E**). Lipids profiling identified major lipid classes and a predominance of phosphatidylcholines (PC) and lysophosphatidylcholines (LPC) and some sphingomyelins (SM) with functional categories related to lysosome and membrane, both involved in EVs secretion (*29*) (**Fig. 1F, fig. S2B**). Together, these results reveal complex RNA, lipid and metabolite signatures in plasma sEVs from adult male mice.

**Fig. 1.**
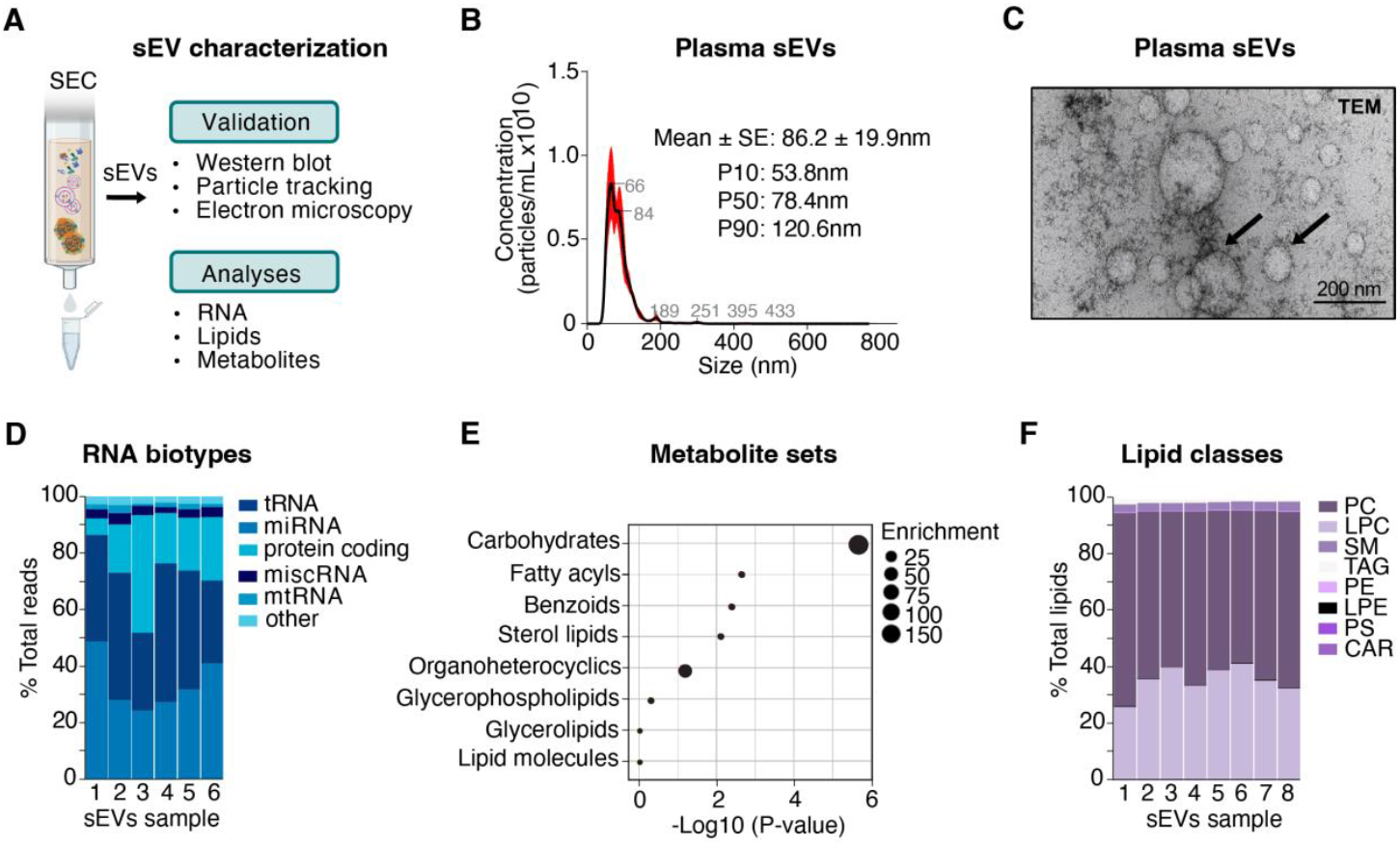
Validation and characterization of mouse plasma sEVs. **A**. Schematic representation of blood plasma sEVs isolation by size-exclusion chromatography (SEC) followed by validation (**B, C, fig. S1**) and analyses (**D-F**). **B, C**. Validation of plasma sEVs from fractions 4-5 (pooled) with concentration of particles (particles/mL) determined by size (nm) obtained by nanoparticle tracking analysis. Mean size (black) +/-standard error (SE, red). P10, P50 and P90 are the mean diameter of particles corresponding to the 10th, 50th and 90th percentile of the sample respectively, and (**C**) representative image of plasma sEVs observed by transmission electron microscopy (TEM). Arrows indicate individual sEVs of different size (range 50-120nm). **D-F**. Analyses of sEVs from fractions 4-5 with (**D**) distribution of RNA biotypes identified by sRNA sequencing. tRNA: transfer RNA, miRNA: microRNA, miscRNA: miscellaneous RNA, mtRNA: mitochondrial RNA. n=6 biologically independent sEVs samples, each sample from an individual mouse, (**E**) enriched sets of metabolites determined by time-of-flight mass spectrometry (TOF-MS), and (**F)**distribution of lipid classes determined by liquid chromatography coupled with mass spectrometry (LC-MS). PC: phosphatidylcholine, LPC: lysophosphatidylcholine, SM: sphingomyelin, TAG: triglycerides, PE: phosphatidylethanolamine, LPE: lysophosphatidylethanolamine, PS: phosphatidylserine, CAR: acylcarnitine. n=8 biologically independent sEVs samples, each sample from an individual mouse. Samples from the same mice were used for RNA, lipid and metabolite analyses.

### The composition of circulating sEVs is altered by ELS

Components of circulating sEVs in blood are influenced by physiological states and can be altered by pathological conditions (*10, 30, 31*). We examined if circulating sEVs are modified by ELS, a condition known to affect plasma composition in humans and rodents (*32*–*34*). We used an established mouse model of chronic stress in early postnatal life (MSUS) (*25*) that induces behavioral, physiological and metabolic phenotypes in exposed animals and their offspring (*24, 25, 35*) (**Fig. 2A**). In males, ELS significantly increased the total number of plasma sEVs without affecting their size (**Fig. 2B, C**). It also altered their RNA, metabolites and lipids content (**Fig. 2D-I**). Thus, small RNAs (sRNAs) including messenger RNA (mRNA)-derived sRNAs, miRNAs and tRNAs from let-7, miR-30 and tRNA-Lys families known to be altered by environmental exposure (*15, 36*–*38*) were down- or up-regulated in plasma sEVs (**Fig. 2D-E, table S1.1**). Specific metabolite sets such as monosaccharides, fatty acids and conjugates including steroid conjugates and bile acids were significantly enriched (**Fig. 2F-G**) and lipids including PCs, PE and TAG were upregulated (**Fig. 2H-I**) in plasma sEVs of ELS-exposed adult males (**table S1.2 and S1.3)**. The overall distribution of lipid classes and functional categories was however unchanged (**fig. S3**).

**Fig. 2.**
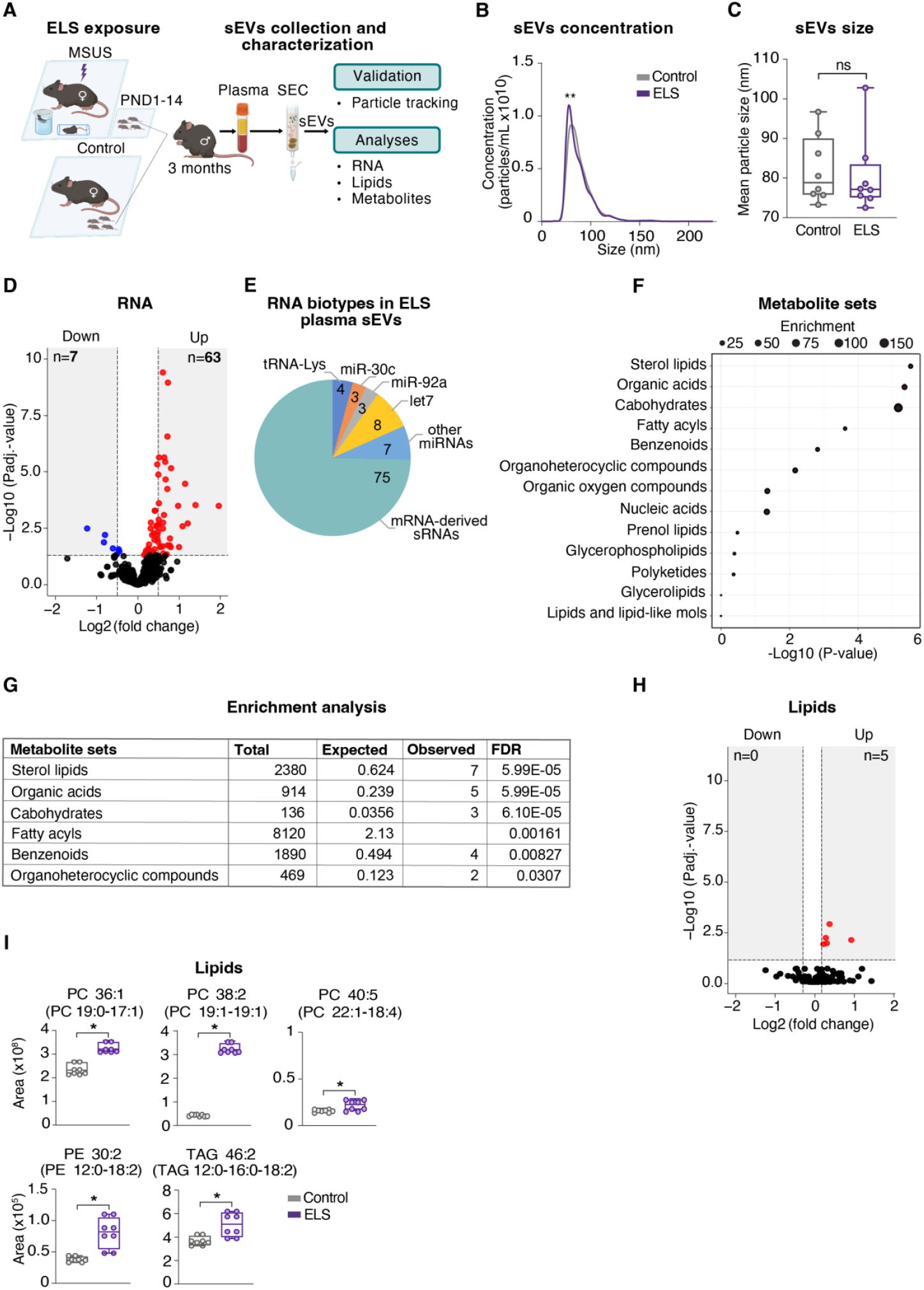
Effects of ELS on RNA, metabolite and lipid content of plasma sEVs. **A**. Experimental design of ELS exposure followed by sEVs collection and analysis. Mouse pups are exposed to unpredictable maternal separation combined with unpredictable maternal stress (MSUS) for 3h daily between postnatal day (PND) 1 and 14. During separation, dams are stressed unpredictably by either 5-min forced swim or 20-min restraint applied randomly. Control animals are left undisturbed. Blood is collected from control and ELS-exposed males when adult (3 months) and plasma sEVs are isolated by SEC for subsequent RNA, metabolite, and lipid, analyses. **B**. Concentration of sEVs in plasma of control and ELS-exposed mice. *n=*8 biologically independent samples per group, each sample from an individual mouse. Mann-Whitney U test, **P<0.01. **C**. Boxplots representing mean particle size of sEVs in plasma of control and ELS-exposed mice. *n=*8 biologically independent samples per group, each sample from an individual mouse. Two-tailed Student’s t-test, ns=non-significant. **D**. Volcano plot of significantly downregulated (blue) and upregulated (red) RNA in plasma sEVs from ELS-exposed males compared to controls. *n=*6 biologically independent samples per group. Benjamini-Hochberg test, P_adj._<0.05. **E**. Pie chart showing the distribution of differentially expressed RNAs based on panel (**D**) in plasma sEVs from ELS-exposed mice. **F**. Enriched sets of metabolites in plasma sEVs from ELS-exposed mice compared to controls. *n=*8 biologically independent sEVs samples per group, each sample from an individual mouse. **G**. List of significantly enriched metabolite sets with total number and enrichment ratio shown as expected versus observed hits with FDR. **H**. Volcano plot of significantly downregulated (blue) and upregulated (red) lipids in plasma sEVs from ELS-exposed males compared to controls. *n=*8 per group for lipids, each sample from an independent mouse. Benjamini-Hochberg test, P_adj._<0.05. **I**. Differentially altered lipids in plasma sEVs from ELS-exposed mice compared to controls. Lipid nomenclature: Number of carbons:Number of double-bonds in fatty acid chain. PC: phosphatidylcholine, PE: phosphatidylethanolamine, TAG: triglycerides. Mann-Whitney U test. Error bars represent SEM, *P<0.05.

We confirmed these results by isolating plasma sEVs using density-gradient ultracentrifugation (DGUC), another method based on density (**fig. S4A**). We identified particles with typical markers and size of sEVs in 2 fractions out of 9 (5-7) **(fig. S4B-D**). In ELS-exposed males (**fig. S5A**), plasma sEVs were significantly more numerous than in controls and contained more miRNAs from let-7 and miR-30 families (**fig. S5B-D**), consistent with SEC results. They had no change in size but notably, DGUC sEVs were slightly larger than SEC sEVs likely due to the different separation method (*39*). Since SEC had a better sEVs yield (10^10^ versus 10^8^ particles/mL plasma) (**Fig. 1B, fig. S4C**), we used it to isolate plasma sEVs for all following experiments.

### Chronic injection of plasma sEVs from ELS-exposed mice alters the metabolic state of naïve mice

Circulating sEVs are associated with organ physiology and not only is their content influenced by cellular states, but they can also themselves modulate cellular pathways involved in e.g. glucose metabolism (*40*). We tested if plasma sEVs from adult males exposed to ELS can modify metabolic functions *in vivo* by injecting them intravenously (i.v.) into naïve adult males (**Fig. 3A**). Plasma sEVs from ELS males (ELS-sEVs) did not change body weight one (D1) or 46 (D46) days after the last injection compared to sEVs from control males but transiently increased weight slightly at D12 (**Fig. 3B**). At D46, the metabolomic profile of plasma was altered in males injected with ELS-sEVs and 18% of all detected metabolites were significantly affected (**Fig. 3C, table S2**). Plasma of males injected with ELS-sEVs had a 100 to 300-fold enrichment in several compounds at D46 including fatty acids, prenol lipids, steroid conjugates and glycerophosphoethanolamines (GPEA) compared to controls **(Fig. 3D)** as well as significant enrichment of pathways involving alpha-linolenic acid and linoleic acid (**Fig. 3E**).Compound analyses of significantly altered metabolites indicated that the differences mostly originate from down-regulated metabolite sets (**fig. S6)**. Notably, the effects ELS-sEVs injection are similar to those observed in males directly exposed to ELS (*34*). When comparing all enriched metabolic pathways and metabolites altered in males injected with ELS-sEVs or males directly exposed to ELS, we observed that changes did not overlap but were unique to each condition (**Fig. 3F**). At D46, plasma lipids were also affected in males injected with ELS-sEVs with <5% of lipids significantly down-regulated and about 50% up-regulated (**Fig. 3g)**. Most significantly increased lipids were PEs and PCs (**Fig. 3H**), similarly to plasma sEVs from ELS-exposed mice (**Fig. 2I**). These results suggest that chronic injection of circulating sEVs from ELS males induces metabolic changes that partially overlap with changes in males directly exposed to ELS and persist for up to 46 days after injection.

**Fig. 3.**
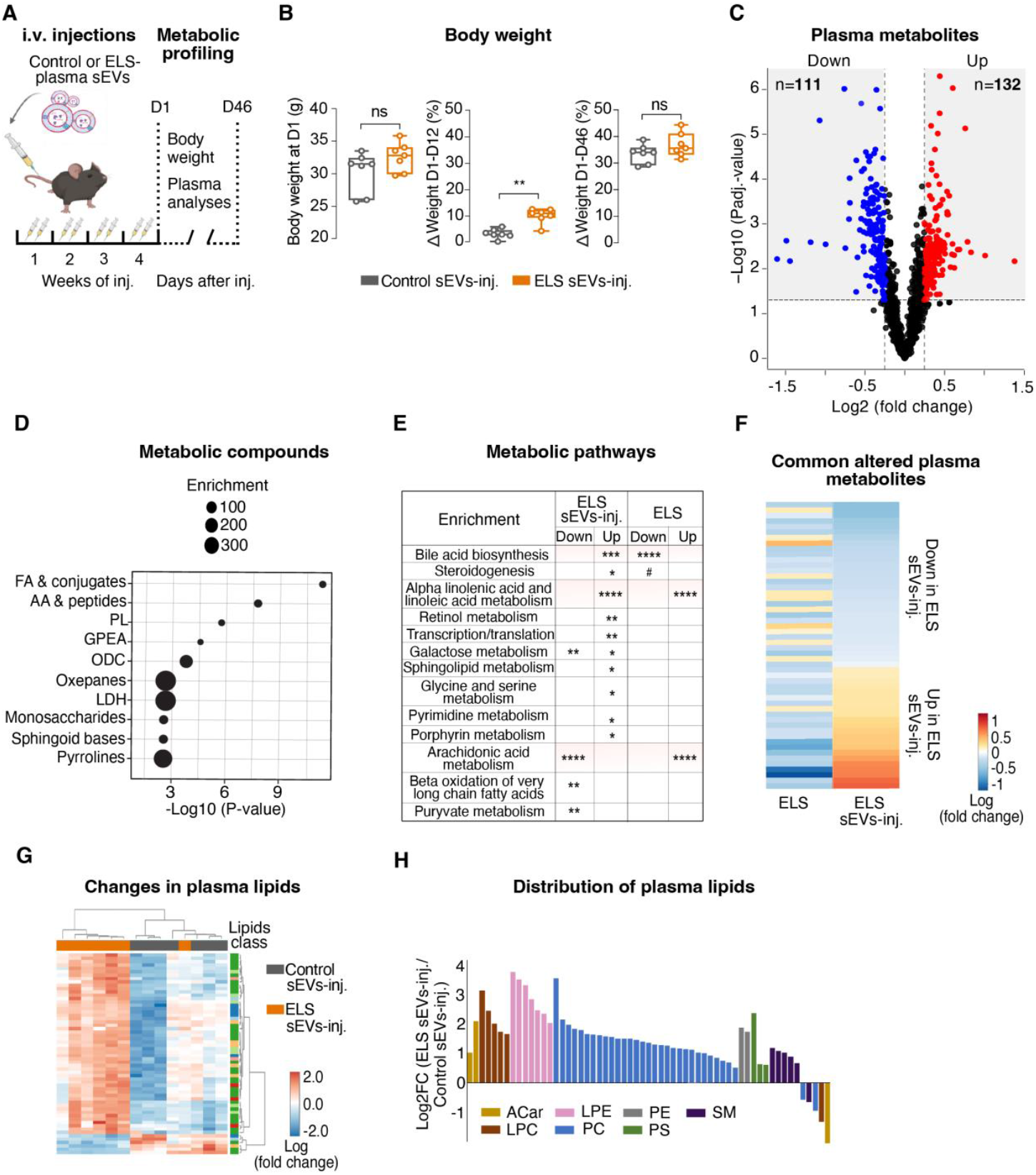
Effects of chronic injection of plasma sEVs from ELS-exposed males on metabolism of naïve mice. **A**. Experimental design for intravenous (i.v.) injection of plasma sEVs and metabolic profiling of injected mice. Plasma sEVs from control or ELS-exposed males are injected into age-matched naïve males twice per week for 4 weeks. Metabolic profiling includes body weight measurement and plasma metabolomic and lipidomic analyses. *n*=7 mice per group. The same animals were used for all analyses in panels (**B-H**). **B**. Body weight of male mice injected with sEVs from control or ELS-exposed males 1 day (D1) after the last injection (left) and % change in body weight between 1 and 12 days (D12, middle) and between 1 and 46 days (D46, right) after injection. Two-tailed unpaired Student’s t-test, ns=non-significant. **P<0.01. **C**. Volcano plot of significantly downregulated (n=111, blue) and upregulated (n=132, red) plasma metabolites in males injected with plasma sEVs from ELS-exposed males at D46. Benjamini-Hochberg test, P_adj._<0.05. Significantly changed metabolites with P_adj._<0.05 were selected as input to assess compound enrichment with MetaboAnalyst tool shown in (**E**). **D**. Metabolic compounds analyses (MetaboAnalyst tool) showing enrichment in plasma of mice injected with sEVs from ELS-exposed mice at D46. FA: fatty acids, AA: amino acids, PL: prenol lipids, GPEA: glycerophosphoethanolamines, ODC: organic dicarboxylic compounds, LDH: linear diarylheptanoids. **E**. Enrichment analysis of metabolites in plasma from ELS-exposed males (our previous data (*34*)) and from males injected with sEVs from ELS-exposed males at D46 (*n*=5 mice per group for controls and ELS; *n*=7 mice per group for injected groups). Benjamini–Hochberg test, ^#^P_adj._<0.1, *P_adj._<0.05, **P_adj._<0.01, ***P_adj._<0.001, ****P_adj._<0.0001. **F**. Heatmap of commonly altered plasma metabolites in ELS-exposed males (ELS (*34*)) and in males injected with plasma sEVs from ELS-exposed males (ELS sEVs-inj) at D46 compared to their respective controls. Benjamini-Hochberg test, P_adj._<0.05. **G**. Heatmap of significantly altered plasma lipids in males injected with plasma sEVs from ELS-exposed males at D46. Benjamini-Hochberg test, P_adj._<0.05. **H**. Distribution of lipid classes of significantly altered lipids in males injected with plasma sEVs from ELS-exposed males at D46. ACar: acylcarnitine, LPE: lysophosphatidylethanolamine, PE: phosphatidylethanolamine, SM: sphingomyelins, LPC: lysophosphatidylcholines, PC: phosphatidylcholines, PS: phosphatidylserine. Benjamini-Hochberg test, P_adj._<0.05.

### Plasma sEVs from ELS-exposed males can signal to the male germline

We previously showed that chronic i.v. injection of plasma from ELS-exposed males induces metabolic phenotypes in the offspring, suggesting germline-dependent intergenerational effects (*34*). We tested if sEVs injected i.v. can reach testis using labeled sEVs. Fluorescent signal was detected in testis of males injected with sEVs tagged with GFP or a lipophilic dye (PKH26) (**Fig. 4A, fig. S7A**). The uptake of labeled plasma sEVs by germ cells was confirmed *in vitro* (**fig. S7B**), suggesting that sEVs are taken up by testicular cells *in vivo* and *in vitro* as previously observed (*41*).

**Fig. 4.**
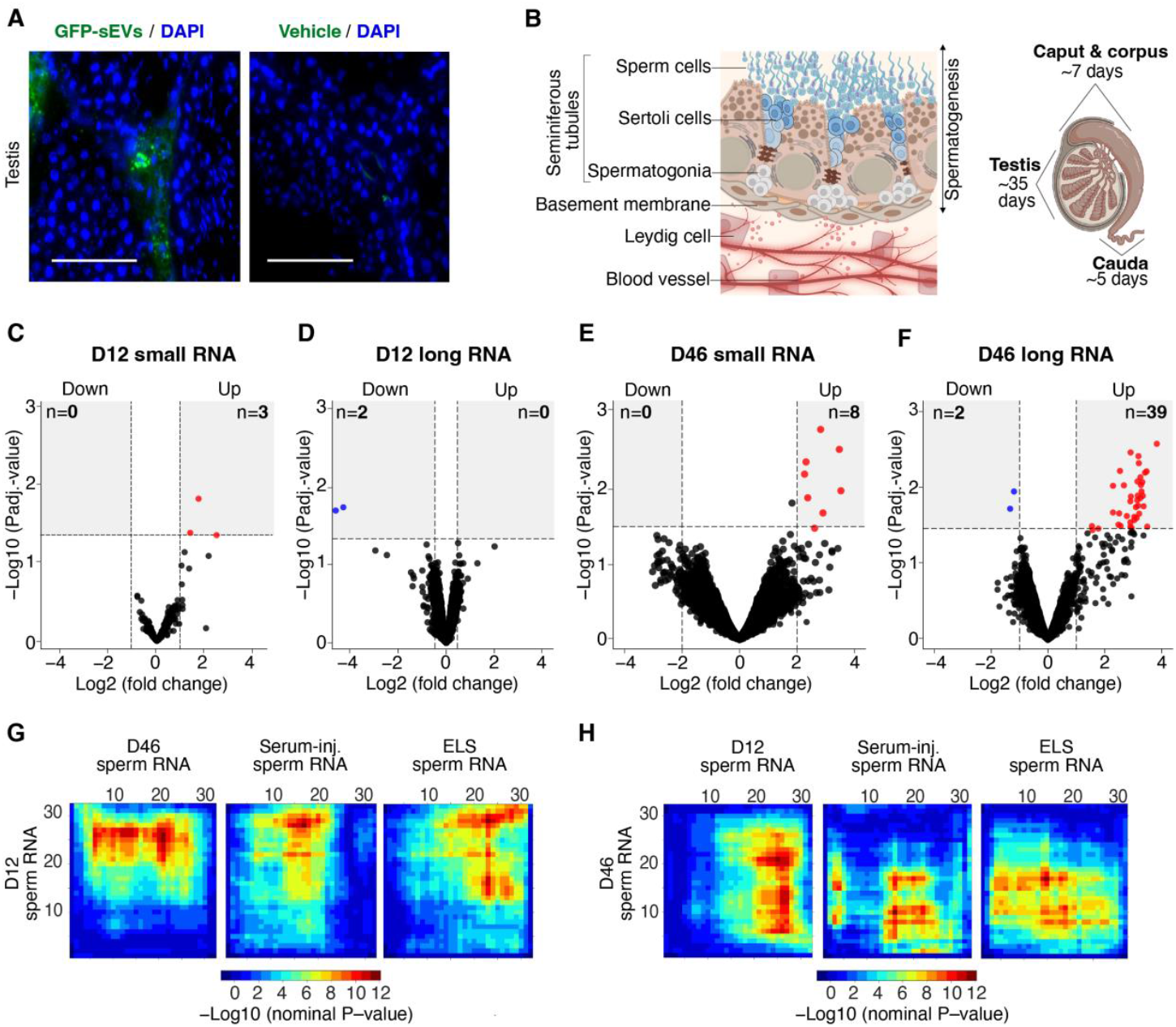
Effects of chronic injection of plasma sEVs from ELS-exposed mice on male germ cells. **A**. Images of testis cross-sections from males injected i.v. with GFP-tagged sEVs and vehicle as control. Nuclei are stained with DAPI. Scale bar: 100μm. **B**. Left: Cross section of a testicular seminiferous tubule showing the different stages of spermatogenesis. In the tubule, spermatogonial cells (germ stem cells) differentiate into spermatogenic cells (blue) that mature into sperm cells released in the lumen of the tube (top). The seminiferous tubule is underlaid by a basement membrane and an interstitial space with Leydig cells and blood vessels. Right: Schematic of a mouse testis and epididymis indicating the time (days) of germ cells development in testis and sperm transit in epididymis. In testis, maturation from spermatogonia to sperm takes **∼**35 days then sperm cells released in epididymis transit through the caput and corpus for **∼**7 days and are stored in cauda for **∼**5 days before spontaneous release or mating. **C-F**. Volcano plots of significantly downregulated (blue) and upregulated (red) small RNAs (left) and long RNAs (right) in sperm from males injected with plasma sEVs from ELS-exposed mice compared to males injected with control plasma sEVs at D12 (**C, D**) and D46 (**E, F**) post-injection. *n*=5 mice per group at D12, *n*=7 mice per group at D46. Benjamini-Hochberg test, P_adj._<0.05. **G, H**. Rank-rank hypergeometric overlap plots for comparison of RNA expression between (**G**) sperm from males injected with plasma sEVs from ELS-exposed mice at D12 and left) sperm from males injected with plasma sEVs from ELS-exposed mice at D46, middle) sperm from males injected with ELS-exposed serum (D12) and right), sperm from ELS-exposed males, and between (**H**) sperm from males injected with plasma sEVs from ELS-exposed mice at D46 and left) sperm from males injected with plasma sEVs from ELS-exposed mice at D12, middle) sperm from males injected with ELS-exposed serum (D12) and right), sperm from ELS-exposed males. Expression levels are represented by -Log10 (nominal P-value) of overlapping RNAs between compared groups.

We next examined the effects of plasma sEVs injection on sperm by collecting sperm cells from injected males at D12 and D46 post-injection. D12 and D46 are key time-points because they allow distinguishing transient effects from spermatogenic cells present in testis at the time of injection from permanent effects derived from germline stem cells (**Fig. 4B**). At D12, sperm cells present in epididymis during injection have been eliminated, thus the collected sperm only originates from testicular spermatogenic cells. Instead at D46, a complete cycle of spermatogenesis has occurred, thus spermatogenic cells present in testis during injection have differentiated into sperm cells and have been eliminated. Collected sperm at D46 then only derives from spermatogonia cells (**Fig. 4B**) (*42*–*44*). At D12 and D46, several small and long RNAs were significantly altered in sperm of males injected with ELS-sEVs with a larger number at D46 (**Fig. 4C-F, fig. S8, table S3 and S4**). Notably, a comparison with our previous data on sperm from ELS-exposed males showed that similar miRNAs are up-regulated in sperm of ELS-sEVs injected males (**table S3.1 and S4.1**) (*20*). These miRNAs have previously been implicated in intergenerational effects of paternal exposure (*15, 36, 37*). Analysis of long RNAs loci showed a significant upregulation of transcription factor (TF) binding sites (340 sites), particularly PPARα-RXRα binding motif that we previously identified in sperm of ELS-exposed adult males (*34*), suggesting functional relevance **(table S3.3)**.

Further analyses of long RNAs in sperm from ELS-sEVs injected males at D12 and D46 using rank-rank hypergeometric overlap (RRHO) revealed two distinct sets of long RNAs: one with increased expression at both D12 and D46 and one with increased expression at D12 but decreased expression at D46 (**Fig. 4G-H**). Further comparative RRHO analyses with our previous data identified common transcripts between ELS-sEVs injected males, males injected with serum from ELS-exposed males (ELS-serum) (*34*) and males directly exposed to ELS (*45*). At D12, sperm from ELS-sEVs injected males had RNA expression patterns similar to sperm from males injected with ELS-serum or males directly exposed to ELS (**Fig. 4G**). In contrast at D46, sperm from ELS-sEVs injected males had mostly opposite expression patterns compared to males injected with ELS-serum or to ELS-exposed males themselves (**Fig. 4H**).

### Plasma sEVs from ELS-exposed mice have intergenerational effects

Our previous work showed that sperm RNA contributes to the transfer of ELS effects from father to offspring (*20, 45*). Since plasma sEVs from ELS-exposed males modify sperm RNA content (**Fig. 4C-F**), we examined if this has phenotypic consequences for the offspring. We bred males injected with plasma ELS-sEVs or control sEVs at D12 and D46, and generated D12 and D46 offspring from these males (**Fig. 5A**). D12 offspring of ELS-sEVs injected males had a slightly lower body weight, lean mass, body fat and water compared to offspring of control sEVs injected males but not D46 offspring, suggesting a transient effect of ELS-sEVs at D12 (**Fig. 5B-C**). However in plasma, metabolic pathways involved in bile acid biosynthesis and/or steroidogenesis were down-regulated in both D12 and D46 offspring of ELS-sEVs injected males, similarly to males directly exposed to ELS and their offspring (*34*) (**Fig. 5D**). Plasma metabolites and related pathways were overall altered in the same direction in D12 and D46 offspring, but metabolites were segregated between the groups, an effect confirmed by a significant number of differentially-expressed metabolites between D12 and D46 offspring of ELS-sEVs injected males (**Fig. 5E-F**). D12 and D46 offspring of ELS-sEVs injected males also had mild alterations in circulating lipids compared to controls (**Fig. 5G**).

**Fig. 5.**
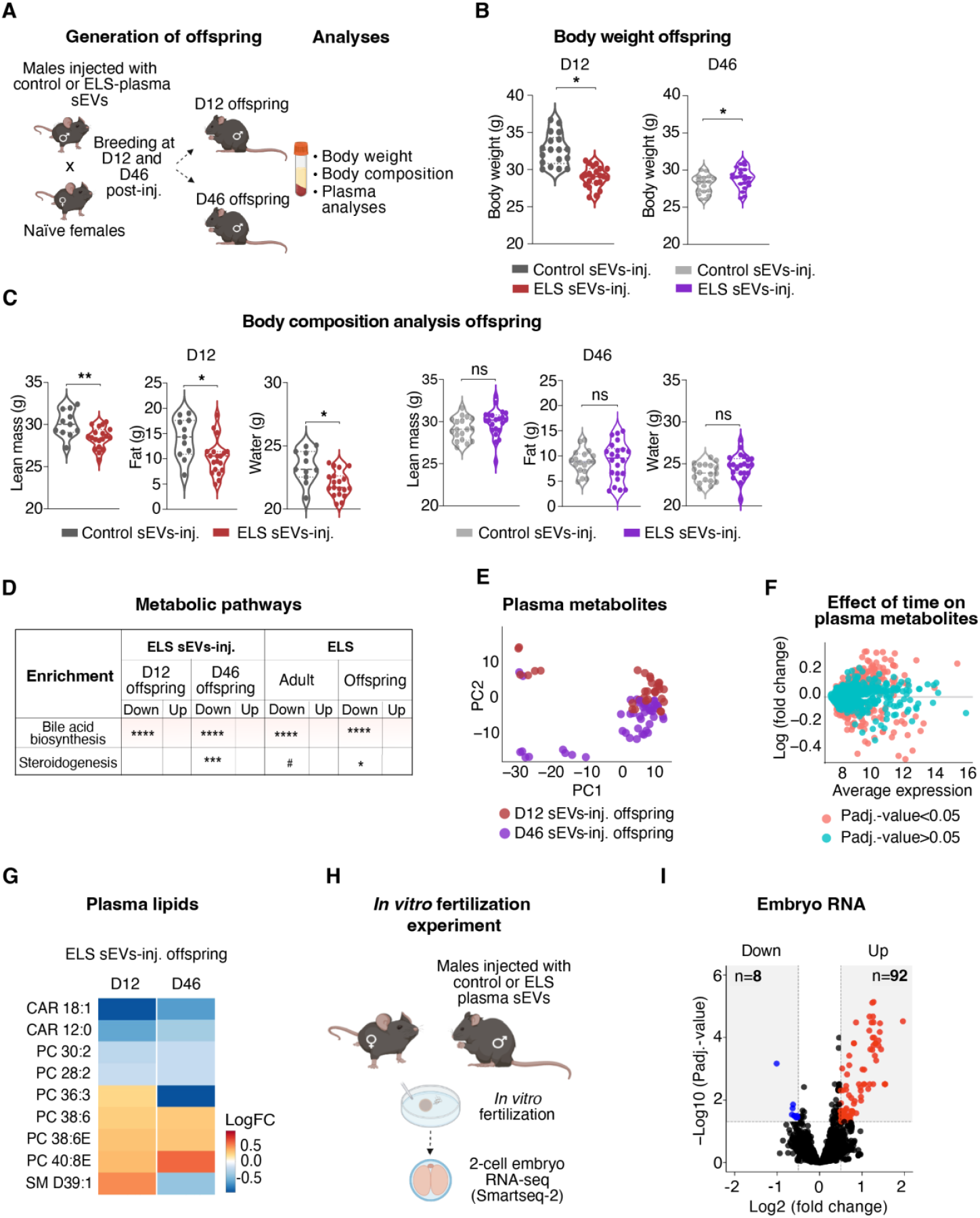
Metabolic changes in the offspring of males injected with plasma sEVs from ELS-exposed males. **A**. Experimental setup for the generation of offspring from males injected with plasma sEVs from control or ELS-exposed males. Males are paired with naïve females at D12 and D46 post-injection and the adult offspring (3-month) is analysed for body weight, body composition by echoMRI and plasma metabolomics. **B, C**. Body weight (**B**) and body composition **(C)** of D12 (left) and D46 (right) offspring of males injected with plasma sEVs from control or ELS-exposed mice. D12: ELS *n*=22, control *n*=18. D46: ELS *n*=22, control *n*=22. Two-tailed Student’s t-test, treatment effect *P<0.05, **P<0.01, ns=non-significant. **D**. Enriched metabolic pathways in D12 and D46 offspring of males injected with plasma sEVs from ELS-exposed mice (left) and in adult ELS males and their adult offspring (right). Statistical analysis done between each experimental group and its corresponding control. Benjamini-Hochberg test, ^#^P_adj._<0.1, *P_adj._<0.05, ***P_adj._<0.001, ****P_adj._<0.0001, ns=non-significant. **E**. Cluster analysis of principal component for plasma metabolites of D12 and D46 offspring of males injected with plasma sEVs from control or ELS-exposed mice. *n*=40 mice at D12, *n*=44 mice at D46. **F**. Effect of time post-injection on plasma metabolite profiles comparing D12 and D46 offspring across control and ELS-exposed groups. *n=*40 mice at D12, *n*=44 mice at D46. **G**. Heatmap of plasma lipids altered in both D12 and D46 offspring of males injected with plasma sEVs from ELS-exposed males compared to D12 and D46 offspring of males injected with plasma sEVs from control males. CAR: Acylcarnitine, PC: Phosphatidylcholine, SM: Sphingomyelin. Benjamini-Hochberg test, P_adj._<0.05. **H**. Experimental setup: Oocytes from naïve females were fertilized *in vitro* with sperm from males injected with plasma sEVs from control or ELS-exposed mice at D12 post-injection. The resulting 2-cell embryos are collected and used for single-cell RNA-sequencing by Smartseq-2. **i**. Volcano plots of significantly downregulated (blue) and upregulated (red) RNAs in 2-cell embryos obtained by IVF from sperm of males injected with ELS-plasma sEVs compared to embryos obtained from sperm of males injected with control plasma sEVs. Benjamini-Hochberg test, P_adj._<0.05.

We examined if the effects are specific to plasma sEVs by repeating the injections using the other plasma components, particularly proteins and lipoproteins (PLP) contained in fractions 8-13 (**fig. S1**). Plasma PLP from ELS-exposed mice had no effect on body weight in injected males at D1 or D46 but led to a transient decrease in weight at D12 and altered circulating metabolites and lipids at D46 (**fig. S9A-C**). However, the offspring at D12 and D46 had normal body weight, suggesting only transient effects of plasma PLP that are not intergenerational (**fig. S9D**).

We next assessed the influence of sperm from sEVs-injected males on the early development of their offspring by generating embryos by *in vitro* fertilization (IVF) (**Fig. 5H**). 2-cell embryos derived from sperm of ELS-sEVs injected males had 100 differentially-expressed RNAs, 90% of which were up-regulated **(Fig. 5I)** and enriched for binding motifs for TFs of zinc-finger (Zf), helix-loop-helix (HLH) and E2F family (**fig. S10A-B, table S5**). Gene ontology (GO) enrichment of differentially-expressed genes showed top terms related to translational functions and RNA splicing processes. Several transposable elements (TEs) were also identified as significantly altered (**fig. S10C-D, table S5.2**). Notably, comparative analyses of transcripts in embryos with miRNAs in sperm used for IVF to generate these embryos (*46*) showed that most targets of up-regulated sperm miRNAs tended to be decreased in embryos, suggesting a functional link between embryo mRNA and sperm miRNAs (**fig. S11**). These results suggest an effect of ELS-sEVs on the next generation via sperm of injected fathers.

## Discussion

This study provides evidence that circulating sEVs are important players in the effects of ELS, carrying molecular signals of stress that can affect the germline and consequentially influence phenotypes in the progeny. By characterizing the RNA, lipid and metabolite content of plasma sEVs in adult male mice exposed to ELS, we show that circulating sEVs are durably modified by such early stress. When chronically injected into naïve males, the altered sEVs can alter the sperm transcriptome as well as embryonic transcriptional programs and metabolic functions in the offspring. The findings highlight the role of circulating sEVs in soma-to-germline communication and provide a potential route for the influence of life experiences on exposed individuals and their descendants.

Circulating sEVs in blood have been reported to be modified by life conditions such as chronic stress (*47, 48*). Our results that plasma ELS-sEVs injected into the bloodstream modify sperm RNA content, extend these findings by showing that the effects of stress on sEVs go beyond circulation and involve the germline. They also complement previous observations that RNA can be transported from somatic to reproductive tissues (*49*–*51*) by identifying plasma sEVs as vectors of transfer of complex molecular signals. Since several sEVs components including RNA, metabolites and lipids are altered by ELS, multiple factors likely contribute to the transfer of ELS signals to the germline. Notably in sperm, miRNAs that are up-regulated by ELS-sEVs injection overlap with miRNAs up-regulated by direct ELS exposure, e.g. let-7c, miR-10b, miR-10b, miR-183, miR-30 and miR-200 family miRNAs. This suggests a link between circulating sEVs and sperm RNA signatures. Several of these miRNAs were reported to be altered in sperm by other exposures or were implicated in other models of paternal transmission (*16, 36*–*38*). The RNA signature of plasma ELS-sEVs is different from the RNA signature of epididymosomes (our previous data (*52*), EVs from the epididymis that load sperm with nucleic acids, proteins and lipids (*53*). This is likely because these sEVs have a different source and origin, whereby plasma sEVs reflect the global physiological state of the whole organism while epididymosomes reflect the state of epithelial cells in the epididymis. Fluorescence imaging confirmed the presence of injected sEVs in germ cells in testis, but the molecular mechanisms underlying sEVs uptake by germ cells and the subsequent RNA integration into the sperm transcriptome are unclear and require further research.

A key finding is that changes in sperm induced by ELS-sEVs affect the offspring and cause differential RNA expression and metabolic phenotypes. These include dysregulated lipid and bile acid pathways, known to be altered by ELS in individuals directly exposed and in their offspring (*34*). These changes imply a mechanistic link between sperm RNA modifications and phenotypic features in the offspring. Notably, miRNAs and long RNAs associated with transcriptional regulation, e.g. PPARα-RXRα motifs, are altered by ELS in sperm suggesting the possible contribution of transcription factors in this link. The observed effects appear to be specific to sEVs and do not involve other circulating protein or lipid components.

The origin of differences in sperm and embryo signatures 12 and 46 days after ELS-sEVs injection is unknown. Our results suggest that the effects of ELS-sEVs injection are dynamic and change across time causing different outcomes. This may be explained by the turnover of spermatogenic cells and the fact that sperm cells at D12 and D46 originate from respectively, late spermatogenic cells and spermatogonia, which have different molecular signatures (*54, 55*) and may be differently affected by ELS.

Complementing earlier studies on epididymal EVs and their localized effects on sperm maturation, our results show how circulating plasma sEVs can be influenced by a systemic stressor in early life and bypass cellular barriers to reach germ cells. This implies that the germline is more integrated into the physiological state of an organism than previously thought. While our study focuses on male sEVs, it is possible that female sEVs are also affected by ELS and pass ELS effects to the germline (*56*). The modulation of EVs cargo by pathological states such as cancer and metabolic disorders has been documented but our findings newly highlight the functional consequences of circulating sEVs in the context of ELS and its intergenerational effects. Further, they suggest that circulating sEVs may be used as readout of an organism’s state of stress or prior stress exposure. These findings may open future work in human populations to assess whether sEVs are biological markers of ELS usable for diagnostics and treatment management of stress-induced mental and metabolic disorders.

Finally, the discovery that stress-induced alterations in plasma sEVs cargo can reprogram the germline has significant implications for the understanding of the biology of heredity.

## Materials and Methods

### Mice

C57Bl/6J mice (Elevage Janvier, Le Genest Saint Isle, France) were used and were kept under a 12h reverse light/dark cycle in a facility with temperature and humidity control and with access to food (M/R Haltung Extrudat, Provimi Kliba SA, Cat. #3436) and water *ad libitum*. Cages contained wood chip bedding (Lignocel select, J. Rettenmaier & Söhne), paper tissue as nesting material and a plastic house. Experiments were run during the active cycle of the animals in conformity with guidelines of the Cantonal Veterinary Office of Zurich and the Swiss Animal Welfare Act (Tierschutzgesetz) and under license number ZH083/2018 and ZH021/2022. Animals were 3-4 months old, except females used for *in vitro* fertilization that were 2 months old.

### Blood collection

For blood collection, adult males were singly housed overnight with food and water prior to sacrifice. Trunk blood was collected into EDTA-coated tubes (Microvette, Sarst-edt) after decapitation and tubes were gently inverted several times. After 15mins at room temperature, blood was centrifuged at 2’000 × rcf for 10min. Plasma was collected, centrifuged at 2’000 × rcf for 5min to remove all debris then stored at -80°C.

### Plasma sEVs and PLPs isolation

To isolate plasma sEVs by SEC, qEVOriginal columns were used (IZON, SP1). Columns were prepared and ran according to the manufacturer’s instructions. 500µL of plasma were loaded and 13 fractions of 1mL immediately collected. SEC fractions 4-5 were used as fractions enriched in sEVs and fractions 8-13 as enriched in PLPs. To isolate plasma sEVs by DGUC, iodoxanol (OptiPrep) density medium (Sigma-Aldrich) was used to prepare an OptiPrep gradient (40%, 20%, 10%, 5%) by diluting OptiPrep with 0.25M sucrose, 10mM Tris. 1mL of plasma was layered onto the OptiPrep gradient and ultracentrifuged at 100’000 x rcf at 4°C for 18h. Ten fractions were collected from top to bottom and each fraction was diluted with filtered cold PBS then ultracentrifuged at 120’000 x rcf for 2h at 4°C. The supernatant was removed, the final pellet was resuspended in cold PBS and frozen at -80°C. Fractions 5-7 of DGUC were pooled and used as fractions enriched in sEVs.

### Sperm collection

Mature sperm was isolated as previously described (*57*). Briefly, cauda epididymis was dissected out and adipose tissue carefully removed. The epididymis was incised with several cuts, placed into M2 medium (M7167-100mL, Sigma-Aldrich) and incubated for 30min at 37°C. The supernatant containing motile sperm was centrifuged at 2’000 x rcf for 10min, mixed with an equal volume of somatic cell lysis buffer (0.1% sodium dodecyl sulfate and 0.05% Triton X-100 in MilliQ water), centrifuged again at 2’000 x rcf for 10min, then washed twice with PBS. The final sperm pellet was snap-frozen and stored at -80°C for further analysis. For *in vitro* fertilization (IVF), cauda epididymis was placed in a petridish with pre-incubated (30mins, 5%CO_2_, 37°C) Card FertiUP medium (Cosmo Bio LTD, KYD-005-EX) then overlayed with mineral oil (M8410-1L, Sigma-Aldrich). One incision per cauda was made and a small ‘ball’ of sperm was dragged through the oil into the drop of FertiUP.

### Western blotting

25μL of each sample were mixed with 10X RIPA (Cell Signaling Technology), incubated for 5min at 4°C then mixed with a 4X Laemmli Sample Buffer (Bio-Rad Laboratories). The membranes were blocked in 5% SureBlock^TM^ (LubioScience) in Tris-buffered saline with 0.05% Tween-20 (TBS-T) (Sigma-Aldrich) for 1h at 20°C and incubated with primary antibodies overnight at 4°C (anti-HSP70 [1:1000; System Biosciences, <u>EXOAB-Hsp70A-1</u>], anti-CD63 [1:1000; System Biosciences, <u>EXOAB-CD63A-1</u>], anti-ApoA1 [1:10000; Genetex, GTX112692], anti-CD9 [1:1000; System Biosciences, <u>EXOAB-CD9A-1</u>], anti-TSG101 [1:2000; Abcam, ab125011]), anti-CD63 [1:2000; System Biosciences, <u>EXOAB-CD63A-1</u>]. Human exosome lysate (System Biosciences, EXOAB-POS-1) was used as positive control. The membranes were then washed in TBS-T, 3 times for 15min, incubated with an HRP-conjugated secondary antibody (anti-rabbit [1:10000; Santa-Cruz Biotechnology, cs2357], anti-mouse [1:10000; Upstate Biotechnology, 12-349]) for 1h then washed again in TBS-T, 3 times for 15min.

### Nanoparticle tracking analysis

The number and size distribution of plasma particles was measured on Nanosight NS300 (Malvern, UK) at 20°C according to the manufacturer’s guidelines. All samples were diluted to a 1:1000 concentration in filtered (0.22µm) PBS (10010-015, Gibco).

### Electron microscopy imaging

sEVs were visualized by transmission electron microscopy (TEM) using a negative stain method with methylcellulose. Briefly, the carrier grid was glow-discharged in plasma for 10min, washed with a 100µL drop of PBS, incubated in 1% glutaraldehyde (GA) in water for 5min, then washed with water 5 times for 2min. The grid was incubated in 1% uranyl acetate (Uac) for 5min and kept on ice in methylcellulose/UAc solution (900µL methylcellulose 2%, 100µL 3% UAc) until imaging.

### RNA extraction

RNA was extracted from plasma sEVs or sperm with Trizol reagent (Life Technologies, 15596026). Briefly for sEVs, samples from SEC or DGUC were lysed in Trizol followed by phenol-chloroform extraction. The RNA pellet was washed in 75% ethanol, air dried and dissolved in nuclease-free water. For sperm, samples were homogenized in Trizol using steal beads and a tissue lyser (TissueLyser II, Qiagen) followed by phenol-chloroform extraction. The final RNA pellet was washed in 75% ethanol, air dried and dissolved in nuclease-free water. RNA concentration and integrity were checked using 2100 Bioanalyzer (Agilent).

### Preparation of sequencing libraries

For plasma sEVs, sRNA libraries were prepared using Smallseq protocol designed for single-cell or low-input RNA sequencing experiments (*58*) and sequenced with Illumina Novaseq6000 (100bp read length, single-end sequencing, mean depth of 18M reads/sample). All samples were sequenced twice independently. For sperm, sRNA-seq libraries were prepared with 3-5ng of total RNA using Realseq Small RNA sequencing library preparation kit (Somagenics) according to the manufacturer’s instructions and sequenced with Illumina Novaseq6000 (100bp read-length, single-end sequencing, mean depth of 30M reads/sample). Total RNA-seq libraries were prepared with 3-5ng sperm RNA using SMARTer® Stranded Total RNA-Seq Kit v3 - Pico Input Mammalian (Takara Bio USA, #634486) according to the manufacturer’s instructions and sequenced with Illumina Novaseq6000 (150bp read-length, paired-end sequencing). A fragmentation time of 150sec and 12 PCR cycles at the final amplification were used. Purifications were performed using AMPure XP beads (Beckman Coulter Life Sciences, #A63881). For embryo RNA sequencing, each single embryo was directly collected (no RNA extraction) into 4µL of lysis buffer (Triton X-100, nuclease free water, RNasin Plus, biotinylated Oligo-dT, dNTP mix) followed by reverse transcription, template switching, PCR preamplification, fragmentation and PCR purification steps according to Smart-seq2 protocol (*59*) then sequenced with Illumina Novaseq 6000 (100bp read-length, single-read sequencing, 5-10M reads/embryo).

### Metabolomics

Metabolites were extracted using an Hamilton STAR M liquid handling robot as follows: 10µL of samples were aliquoted into 1.2mL 96 deep-well plates and 300µL of cold extraction solvent methanol:H2O (4:1 v:v) were added. After brief vortexing on a plate shaker, samples were kept at

-20°C for 2h then centrifuged at 4’000 x rcf for 20min. The supernatant was transferred to another deep-well plate and dried under N_2_. Before mass spectrometry analysis, dried extracts were resuspended in 100 µL MilliQ H2O, transferred to PCR 96 deep-well plates and sealed using heat. Flow injection time-of-flight mass spectrometry was done as described previously (*60*). Samples were directly injected into the mass spectrometer (Agilent QTOF 6546) using an autosampler (Gerstel) and a quaternary pump Agilent 1100. Samples were injected at an isocratic flow rate of 150µL/min of solvent isopropanol:H2O (6:4, v:v) containing ammonium fluoride (1mM) and the reference compounds hexakis (2,2,3,3-tetrafluoropropoxy phosphazene) and homotaurine (3-amino-1-propane sulfonic acid). Electrospray ionization was used with the following source parameters: gas temperature 225°C, drying gas 5L/min, nebulizer 20psi, sheath gas temperature 350°C, sheath gas flow 10L/min, Vcap 3’500V and nozzle voltage 1’000V with the fragmentor set to 120V, skimmer to 65V and the Oct1RF Vpp to 750V. Mass spectrometry was operated in full scan mode to scan the mass range (50-1000m/z) at 1.4 spectra/sec. Online mass correction was done using the reference masses 138.0230374 and 940.0003763.

### Lipidomics

Lipids were extracted as described previously (*61*) with some modifications. 20µL of samples were mixed with 1mL of methanol:MTBE:chloroform (MMC) 1.33:1:1 (v/v/v), vortexed briefly then continuously mixed in a thermomixer (Eppendorf) at 25°C (950rpm) for 30min. Proteins were precipitated by centrifugation at 16’000g for 10min at 25°C, the single-phase supernatant was collected, dried under N_2_ and stored at -20°C. Prior to analysis, the dried lipids were redissolved in 100µL MeOH:isopropanol (1:1). Liquid chromatography was done as described previously (*62, 63*) with some modifications. A Vanquish LC pump (Thermo Scientific) was used with the following mobile phases: (a) acetonitrile:water (6:4) with 10mM ammonium acetate and 0.1% formic acid, (b) isopropanol:acetonitrile (9:1) with 10mM ammonium acetate and 0.1% formic acid. Lipids were separated with C18 reverse phase chromatography using Acquity BEH column (Waters) of 100mm length, 2.1mm internal diameter and 1.7µm particle diameter. The column temperature was set to 60°C. The following gradient was used with a flow rate of 1.2mL/min: 0.0-0.29min (isocratic 15-30%B), 0.29-0.37min (ramp 30-48%B), 0.37-1.64min (ramp 48-82%B),1.64-1.72min (ramp 82-99%B), 1.72-1.79min (isocratic 99%B), 1.79-1.81min (ramp 100-15%B) and 1.81-2.24min (isocratic 15%B). Injection volume was 2µL. Needle wash solvent was methanol:isopronal:acetontirtile:H2O (1:1:1:1, v:v:v:v). The liquid chromatography was coupled to a hybrid quadrupole-orbitrap mass spectrometer (Q-Exactive HFx, Thermo Scientific). Heated electrospray ionization was used with the following source parameters: sheath gas flow rate 40, aux gas flow rate 8, spray voltage 3.5kV, capillary temperature 300°C, funnel RF level 50, auxiliary gas heater temperature 300°C. Mass spectrometry was operated in data-dependent acquisition mode (DDA). Full scanning from 200-2000m/z at a resolution of 60’000 and AGC Target 1e6, max injection time 100ms was done. The top 2 precursors were automatically selected for fragmentation using normalized collision energies (NCE) of 20, 30,50 at a resolution of 7’500 and an AGC target of 1e5.

### Early life stress (ELS)

ELS was induced using the MSUS paradigm as previously described (25). Briefly, 3-month-old C57Bl/6J females and males were paired for 1 week to obtain offspring. Following delivery, litters were randomly allocated to ELS or control groups with litter size and number of animals balanced between both groups. Dams assigned to ELS group were unpredictably separated from their pups for 3h per day, from postnatal day (PND) 1-14. During separation, dams were subjected to either a 5-min acute forced swim in cold water (18°C) or a 20-min restraint stress in a tube. At PND21, pups from both groups were weaned and assigned to cages of 3-5 animals/cage, based on gender and treatment group. To avoid litter effects, pups from the same litter were distributed to different cages. Control animals were left undisturbed apart from a cage change once a week until weaning at PND21.

### Intravenous injections (sEVs and PLPs)

sEVs and PLP freshly isolated by SEC (100µL, ∼10^10^ sEVs) from adult male mice exposed to ELS and control males were injected intravenously (i.v.) twice per week for 4 weeks into adult naïve males. For injection, each mouse is placed in a restraint tube, its tail warmed in water and either one of the two lateral veins are used in alternation for injection. Successful injection was confirmed by the absence of resistance and no bleeding. Each injected male was paired with two primiparous adult naïve females both 12 and 46 days after the last injection to obtain D12 and D46 offspring. Body weight was measured at D1, D12 and D46. Metabolomic and lipidomic analyses were conducted at D46. For IVF experiments, sEVs injections were conducted using the same protocol using an independent batch of naïve males.

### Body weight and body composition analysis

Body weight was measured with a simple scale. Total body fat and lean and total water mass were measured with an EchoMRI^TM^ system. For this, males were placed and fixed in a plastic holder that was then inserted into a tubular space at the side of the EchoMRI^TM^ system. Each animal was scanned for 2.5-3min.

### sEVs uptake in vivo and in vitro

For *in vivo* localization experiments, GFP-tagged EVs from HEK cells (System Biosciences, XPAK100EX-G), plasma sEVs labeled with PKH26 dye according to the manufacturer’s protocol (Sigma-Aldrich) or vehicle control were injected i.v. into adult males, following which the animals were sacrificed and testes were collected 120min after injection. Testes were fixed in 4% paraformaldehyde (PFA) overnight at 4°C, dehydrated in an ethanol gradient and embedded in paraffin. The tissue was sliced in 5-8µm thick sections for visualization. Nuclei were stained with DAPI and imaging was done using a confocal microscope. For *in vitro* uptake, GC-1 cells (ATCC CRL-2053TM) were cultured at 37°C in Dulbecco’s modified Eagle’s medium (DMEM-high glucose, Sigma-Aldrich) supplemented with 10% (v/v) foetal bovine serum (FBS, HyClone) and 40lg/ml gentamicin (Sigma-Aldrich). 10^5^ cells were seeded per well in a 24-well plate for 24h then cells were incubated with ∼10^10^ plasma sEVs labeled with PKH26 or PKH67 (Sigma-Aldrich). After 3h of incubation, the cells were washed with PBS to remove unbound exosomes, fixed with 4% PFA and stained with DAPI.

### In vitro fertilization (IVF) and embryos collection

Female mice were superovulated by injection of one dose of Card Hyperova (Cosmo Bio, KYD-010-EX-X5) followed by 5U of human chorionic gonadotropin (hCG) 48h later. 16h after hCG injection, the females were sacrificed by cervical dislocation and the ampula excised with scissors. Cumulus-oocyte complexes (COC) were released from the ampula and placed into a drop of human tubal fluid (HTF, Cosmo Bio LTD, CSR-R-B070) overlayed with mineral oil (Sigma,

M8410-1L) and incubated for 1h at 37°C (5%CO_2_). Sperm was incubated in a drop of Card FertiUP (Cosmo Bio LTD, KYD-002-05-EX) overlayed with mineral oil (M8410-1L, Sigma) and allowed to disperse in the medium for 1h at 37°C (5%CO_2_). Sperm was added to the drop of oocytes and the dish was incubated at 37°C for 3h for fertilization. The samples were washed 3 times with HTF medium to remove cell debris, degenerating oocytes and dead sperm. Fertilized oocytes (zygotes) were moved to a fresh drop of HTP, overlayed with oil and incubated at 37°C overnight. The next morning, 2-cell embryos were collected into a tube containing lysis buffer for Smart-seq2 analyses.

### Bioinformatic data analyses

#### Plasma sEVs sRNA sequencing

FASTQ raw read files were processed as previously described (*58*). Briefly, unique molecular identifiers (UMIs) on reads 5’ end were appended on the read name using UMI-tools (*64*) and adapters (CA-linker) were removed from the beginning of each read using Cutadapt (*65*). Reads were mapped to the mouse genome with STAR aligner (*66*) and final read count table was used for differential expression analysis between groups using edgeR (*67*). Batch correction was done to avoid falsely increasing power due to two sequencing runs conducted on all samples.

#### Sperm sRNA-seq and long RNA-seq

Cutadapt was used to trim the sequence TGGAATTCTCGGGTGCCAAGG according to manufacturer’s instructions. Reads were processed by integrating ExceRpt pipeline for sRNAs analyses (*67*). UMI-tools (*64*) was used to trim the 8nt UMIs and Cutadapt to remove 3nt UMI linker and 3nt Pico v3 SMART UMI Adapter from Read2 prior to mapping. Mapping to the mouse genome (mm10) was done with hisat2 tool (*68*) followed by sorting and indexing with samtools (*69*). Final bam files were deduplicated with UMI-tools and raw counts were calculated with featureCounts of the Rsubread package (*70*). Hypergeometric overlap analyses were conducted to assess the statistical overrepresentation of specific RNA species or functional categories among the identified differentially expressed transcripts using the *phyper* function in R without setting arbitrary significance thresholds.

#### Embryo Smart-seq2

Adapters were removed with Cutadapt then reads were mapped to the mouse genome (mm10) with STAR aligner (*66*) following raw count quantification with featureCounts (*71*). Differential expression was assessed in edgeR (*67*). Genes with FDR<0.05 were used for functional enrichment analysis using g:Profiler web-tool (*72*). To differentiate stages of development of 2-cell embryos, gene expression profiles were projected onto an index with profiles from early, mid and late 2-cell embryos (*73*) using SCMAP tool (*74*).

#### Metabolomics

Raw mass spectrometric data were converted to an open source format (mz5) and processed using an in-house data analyses pipeline (FiaMiner) in Matlab. The m/z axis for the whole dataset was recalibrated using expected reference masses. Annotation was done based on m/z, with a mass accuracy of 0.001Da matching the Human Metabolome Database (HMDB). Differential analysis statistics were performed in Matlab for pairwise and group comparisons with correction for multiple comparisons. Pathway enrichment analysis was performed using HMDB pathway definition v 3.0. Metabolite set enrichment analysis was performed with MetaboAnalyst tool by looking at enriched sets of functionally related metabolites (*75*).

#### Lipidomics

Raw mass spectrometric data were imported in Compound Discoverer 3.1 (Thermo Scientific) for data analysis. Peak picking, retention time alignment and compound grouping were performed.

Lipid annotation was done by matching MS2 spectra to LipidBlast in-silico library. Lipid identification was manually confirmed based on lipidomics standards’ initiative criteria. Annotations not matching the stringent criteria were filtered out. Peak areas were normalized using median normalization and differential analysis was conducted between the different groups. Bottom of Form

## Acknowledgments

We thank Kristina Thumfart and Martin Roszkowski for technical help and advice, Pawel Pelczar and his team for teaching and advising on IVF experiments, Dimitri Schmid, Lola Kourouma, Chiara Boscardin and Anastasiia Efimova for help in conducting IVF and animal experiments, Katharina Gapp for advice on IVF and scientific concepts and methods, Alekhya Mazukhar for initial training with nanoparticle tracking-analysis, Emilio Yangüez for advice on RNA-seq experiments, Johannes Riemann for help with EM images, Simon Berger for help with microscopy, Michael Hagemann-Jensen for advice on Smallseq protocol, Kerem Uzel, Deepak Tanwar and Pierre-Luc Germain for advice on data analysis, Emmanuelle Maciel for technical help. We thank Yvonne Zipfel and LASC team for taking care of the animals. We are grateful to Ali Jawaid, Eloise Kremer and Mia Holmes for their help during the initial stages of the project. We thank Nancy Carullo for initial feedback on the manuscript and Ellen Jaspers for help with the final stages of the manuscript.

## Funding

University Zürich, Switzerland (IMM)

ETH Zürich, Switzerland (IMM)

Swiss National Science Foundation grant 31003A_175742/1 (IMM)

National Centre of Competence in Research (NCCR) RNA&Disease, funded by the Swiss National Science Foundation grants 182880/Phase 2 and 205601/Phase 3 (IMM)

ETH grants ETH-10 15-2 and ETH-17 13-2 (IMM)

European Union Horizon 2020 Research Innovation Program grant 848158 (IMM)

European Union projects FAMILY and HappyMums, both funded by the Swiss State Secretariat for Education, Research and Innovation (SERI) (IMM)

FreeNovation grant from Novartis Forschungsstiftung (IMM)

The Escher Family Fund (IMM)

ETH Postdoctoral Fellow grant 20-1 FEL-28 (RGA-M)

## Author contributions

Conceptualization: AA, IMM

Methodology: AA, FM, LCS, IMM

Investigation: AA, NZ, AO

Visualization: AA, RGA-M

Funding acquisition: IMM

Project administration: IMM

Supervision: IMM

Writing – original draft: AA, IMM

Writing – review & editing: IMM, RGA-M, LCS

## Data and materials availability

All datasets in this study will be made publicly available upon acceptance of the manuscript. RNA-seq datasets will be deposited to the National Center for Biotechnology Information Gene Expression Omnibus (NCBI GEO) and metabolomic and lipidomic datasets to MetaboLights.

