## Supplementary material for "Circulating extracellular vesicles can transport stress signals to the male germline": Suppl Figures 1-11

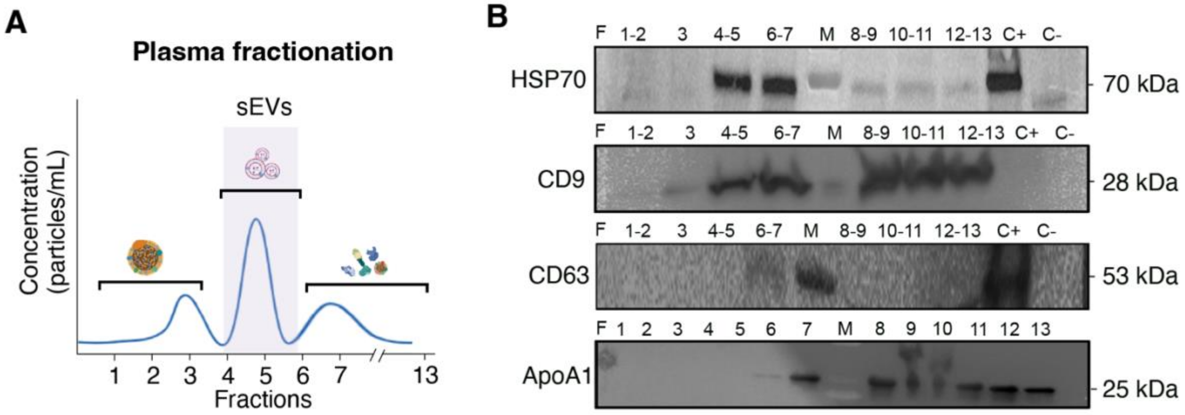


**Fig. S1. Fractionation of mouse blood plasma and analyses by Western blotting. A.** Blood plasma was fractionated by size-exclusion chromatography (SEC) and a total of 13 fractions were obtained. Fractions were separated into three major populations corresponding to large EVs (fractions 1-3), small EVs (sEVs) (fractions 4-6), proteins and lipoproteins (fractions 7-13). **B.** Western blot analysis with common sEVs markers show that fractions 1-3 do not contain any sEVs proteins, fractions 4-5 and 6-7 contain the heat-shock protein and cytosolic marker HSP70 (70kDa) and tetraspanin family protein CD9 (28kDa) but no HDL, and fractions 6-7 contain in addition CD63 (53kDa). Fractions 6-13 contain the HDL marker ApoA1 (25kDa), a known contaminant in plasma sEVs preparations. sEVs fractions 4-5 were selected and pooled for validation and analyses (**Fig. 1**). F: fraction, M: molecular weight marker, C+: positive control (human exosome lysate), C-: negative control (buffer).

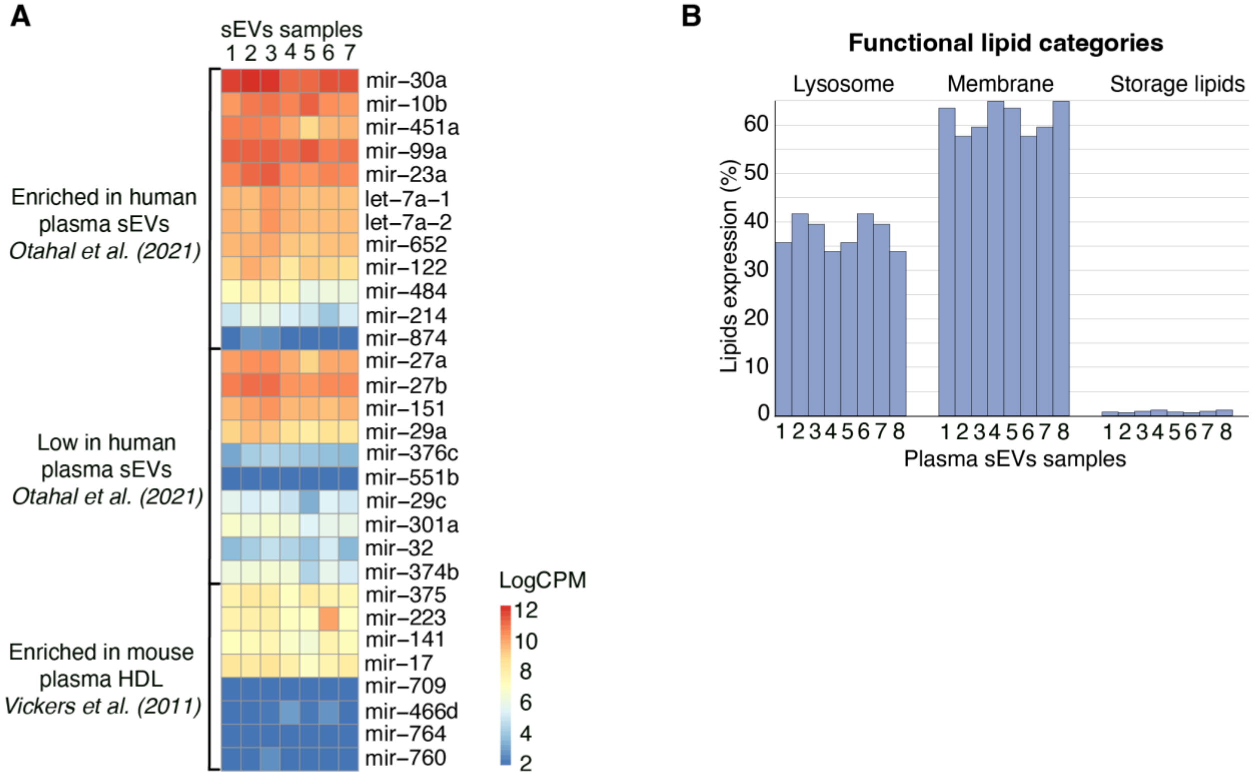


**Fig. S2.** Expression of sEVs-specific and HDL-specific miRNAs and functional lipid categories in plasma sEVs. A. Heatmap showing the level of expression of known sEVs-specific and HDL-specific miRNAs in human and mouse plasma sEVs isolated by SEC previously reported (27, 28). *n*=7 biologically independent sEVs samples, each sample from an individual mouse. B. Analyses of functional categories related to the cellular localization of lipid moieties in mouse plasma sEVs isolated by SEC. *n*=8 biologically independent sEVs samples, each sample from an individual mouse.

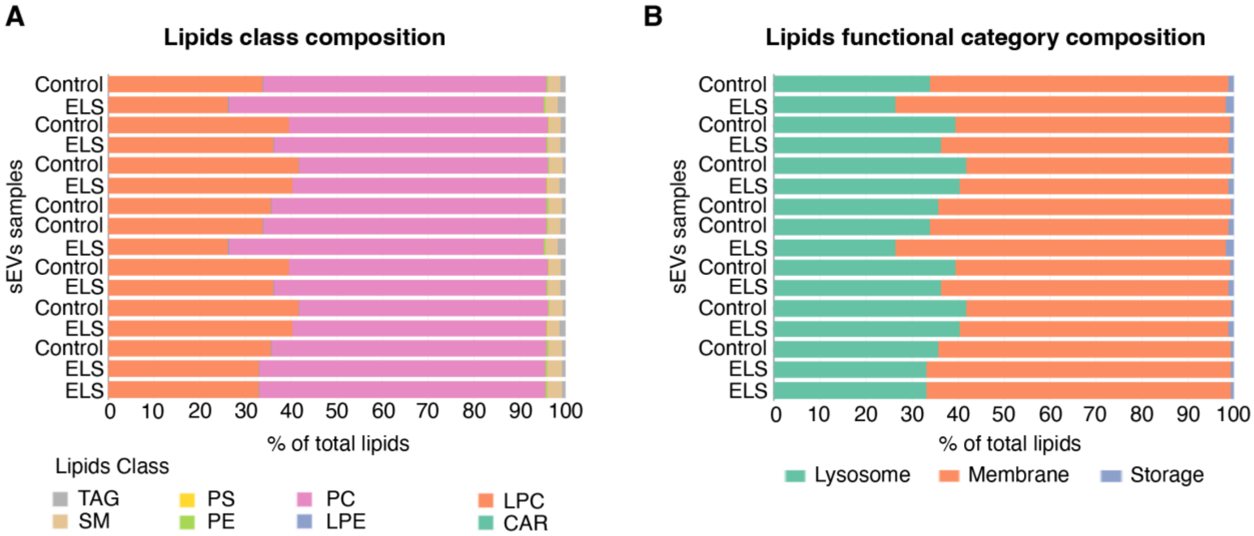


**Fig. S3. Lipid composition and functional categories of plasma sEVs. A, B**. Composition of lipid classes (**A**) and functional categories (**B**) expressed in % total lipids in plasma sEVs from ELS-exposed or control mice. *n*=8 biologically independent samples per group, each sample from an individual mouse. TAG: Triglyceride, SM: Sphingomyelin, PS: Phosphatidylserine, PE: Phosphatidylethanolamine, PC: Phosphatidylcholine, LPE: Lysophosphatidylethanolamine, LPC: Lysophosphatidylcholine, CAR: Acylcarnitine.

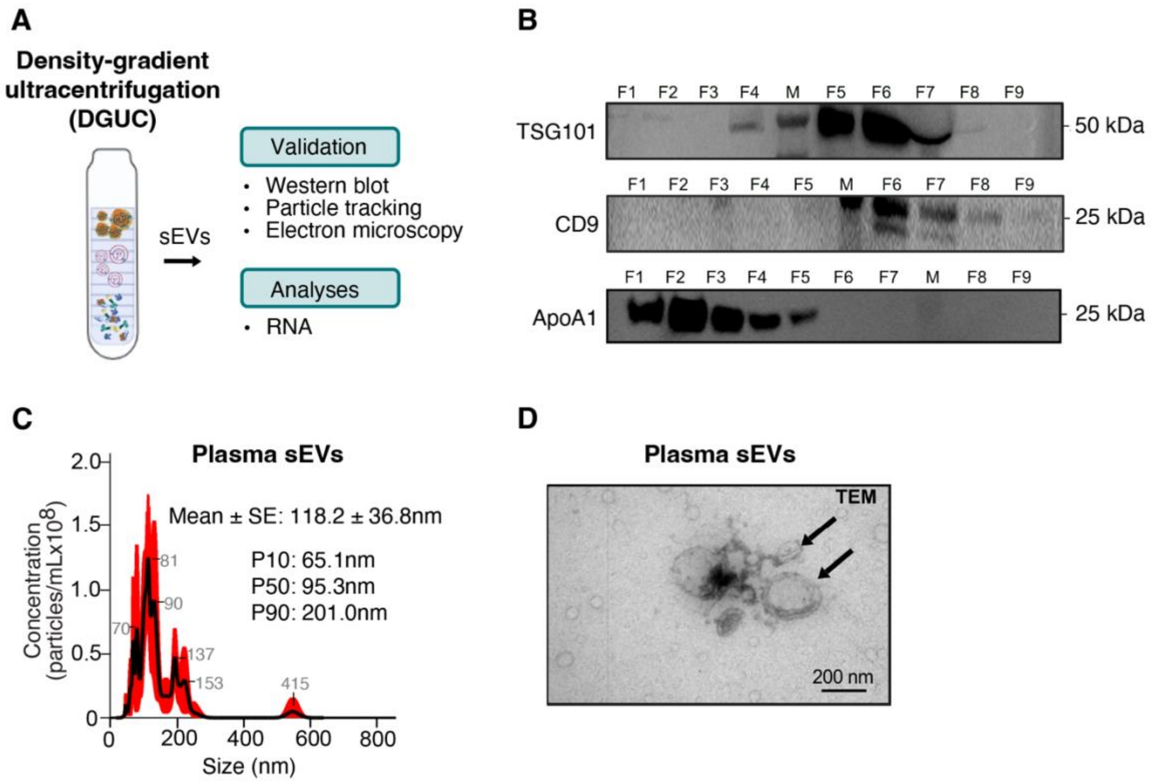


**Fig. S4. Isolation of mouse plasma sEVs by density-gradient ultracentrifugation (DGUC) and characterization. A.** Schematic representation of plasma sEVs collection by DGUC followed by validation and analysis. **B.** Western blot analysis showing the level of sEVs markers including the endosomal sorting complex required for transport (ESCRT)-associated protein TSG101 (50kDa), tetraspanin family protein CD9 (25kDa) and HDL marker ApoA1 (25kDa) in the 9 plasma fractions collected by DGUC. F: fraction, M: molecular weight marker. **C.** Concentration of particles (particles/mL) in plasma fractions 6-8 (F6-F8) determined by size (nm) obtained by nanoparticle tracking analysis. Mean size (black) +/- standard error (SE, red). P10, P50 and P90 represent the mean diameter of particles corresponding to the 10$^{th}$, 50$^{th}$ and 90$^{th}$ percentile of the sample respectively. **D.** Representative image of plasma sEVs isolated by DGUC observed by transmission electron microscopy (TEM). Arrows indicate individual sEVs of different size (range 65-200nm).

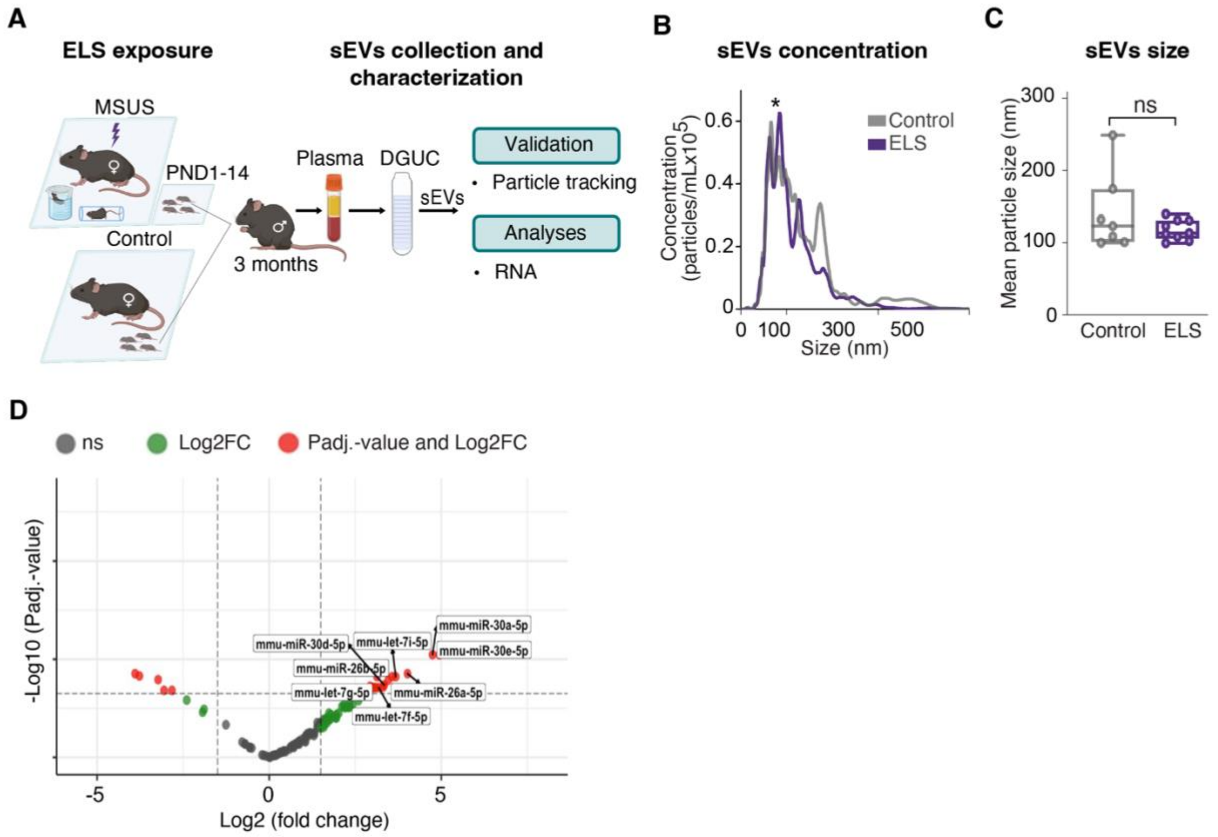


**Fig. S5. Analyses of the effects of ELS on mouse plasma sEVs isolated by DGUC. A.** Experimental design of ELS exposure followed by sEVs collection and analysis. Blood is collected from 3-month-old control and ELS-exposed males, plasma is prepared and fractionated by DGUC to isolate sEVs that are then validated and analysed. The same samples were used for panels (**B-D**). **B.** Concentration of sEVs isolated by DGUC in plasma of control and ELS-exposed males. Mann-Whitney U test, *P<0.05. **C.** Boxplots of mean particle size of sEVs isolated by DGUC from plasma of control or ELS-exposed males. *n*=7 control, *n*=9 ELS samples, each *n* represents a pool of 2 mice. Two-tailed Student's t-test. ns=non-significant. **D.** Volcano plot of differentially expressed small RNAs in plasma sEVs from ELS-exposed males compared to controls. Benjamini-Hochberg test, $P_{adj.}<0.05$.

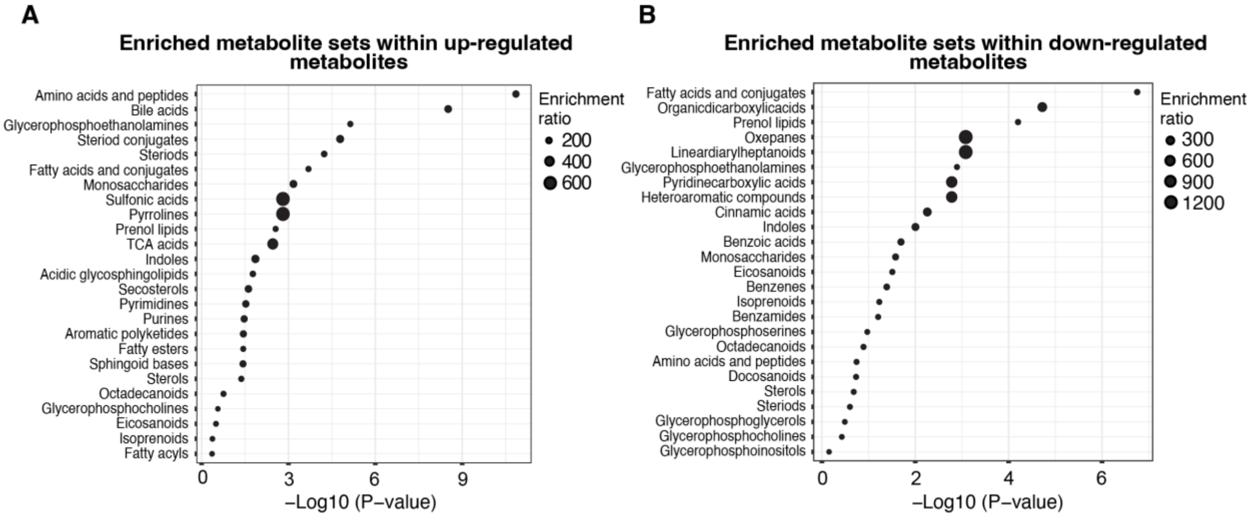


**Fig. S6. Impact of chronic injection of plasma sEVs on metabolism in naïve mice. A, B.** Diagram showing metabolite sets enriched within differentially upregulated (**A**) and downregulated (**B**) individual metabolites in males injected with plasma sEVs from ELS-exposed males compared to controls. *n*=7 biologically independent samples per group, each sample represents an individual mouse. Benjamini-Hochberg test, $P_{adj.}<0.05$.

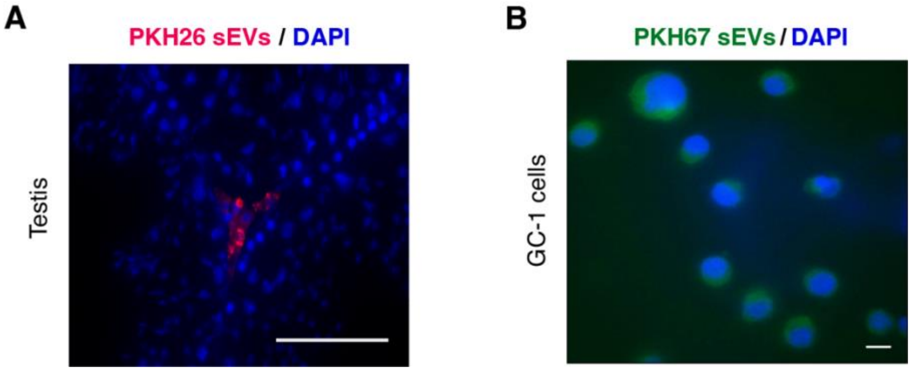


**Fig. S7. Visualization of plasma sEVs *in vivo* and *in vitro*. A.** Cross-section of testis from an adult male 2h after i.v. injection of plasma sEVs labeled with PKH26, a red fluorescent dye, showing signal in testis cells. Nuclei were stained with DAPI. Scale bar: 100μm. **B.** Fluorescence microscopy image of germ cell-like GC-1 cells treated with plasma sEVs labeled with PKH67 (green) dye showing fluorescent signal in cells cytoplasm. Nuclei were stained with DAPI. Scale bar: 10 μm.

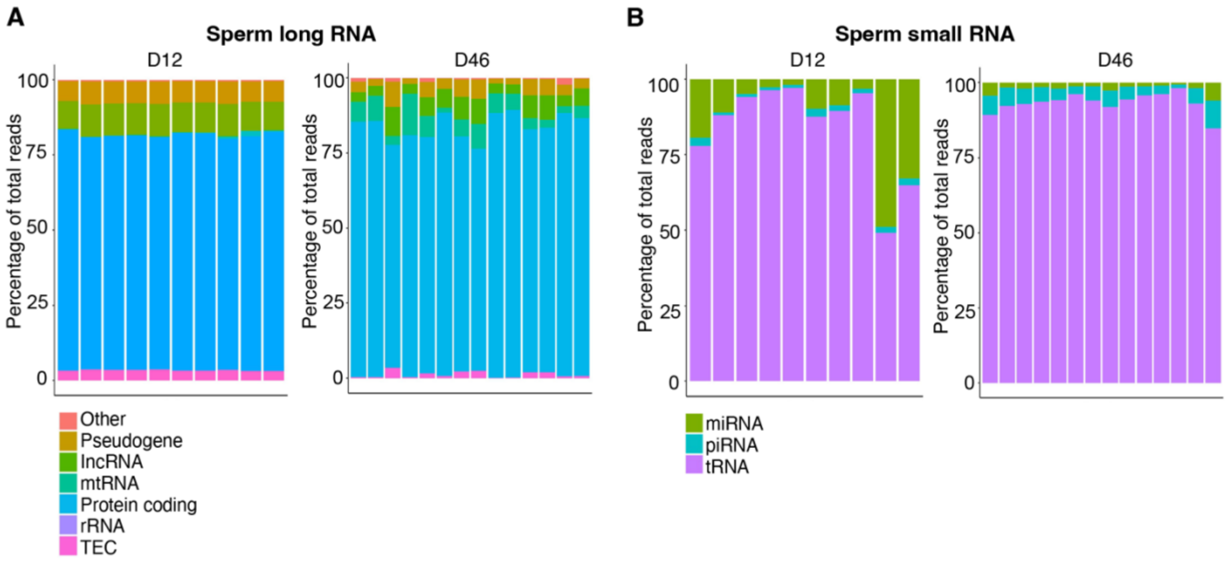


**Fig. S8. Effects of plasma sEVs injection on sperm RNA of injected males.** Distribution of long (**A**) and small (**B**) RNAs biotypes in sperm from males injected with plasma sEVs from control mice and from males injected with plasma sEVs from ELS-exposed mice at D12 (*n*=5 males per group) and D46 (*n*=7 males per group). lncRNA: Long non-coding RNA, mtRNA: Mitochondrial RNA, rRNA: Ribosomal RNA, TEC: Trans-spliced exon coupled RNA, miRNA: Micro-RNA, piRNA: Piwi-interacting RNA, tRNA: Transfer RNA.

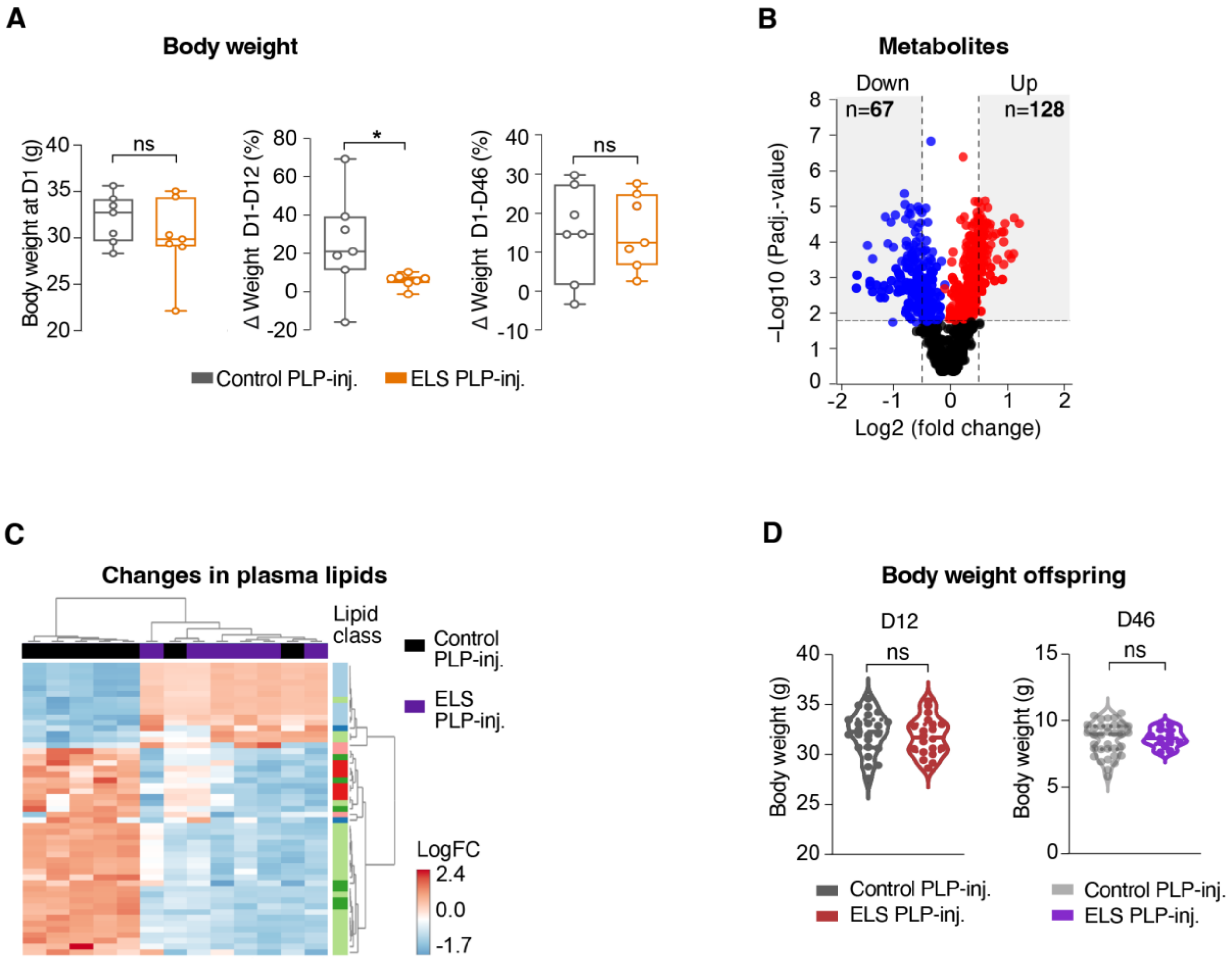


**Fig. S9. Effects of injection of proteins and lipoproteins (PLP) from ELS plasma on metabolism in naïve males and their offspring. A.** Body weight of male mice injected with PLP from control or ELS-exposed males 1 day (D1) after the last injection (left) and % change in body weight between D1-D12 (middle) and D1-D46 (right) post-injection of male mice injected with PLP from control or ELS-exposed plasma. *n*=7 mice per group. Two-tailed unpaired Student's t-test, *P<0.05, ns=non-significant. **B.** Volcano plot of significantly downregulated (n=67, blue) and upregulated (n=128, red) metabolites in plasma of males injected with ELS-exposed PLP at D46 post-injection compared to controls. *n*=7 mice per group. Benjamini-Hochberg test, $P_{adj.}$<0.05. **C.** Heatmap of significantly altered plasma lipids in males injected with ELS-exposed PLP at D46 post-injection compared to controls. *n*=7 mice per group. Benjamini-Hochberg test, $P_{adj.}$<0.05. **D.** Body weight of the offspring of males injected with control or ELS-exposed PLP at D12 and D46 post-injection. *n*=24 offspring from 7 control PLP-injected males, *n*=20 offspring from 6 ELS PLP-injected males at D12. *n*=36 offspring from 7 control PLP-injected males, *n*=13 offspring from 4 ELS-exposed PLP-injected males at D46. Two-tailed unpaired Student's t-test, *ns*=non-significant.

**A**

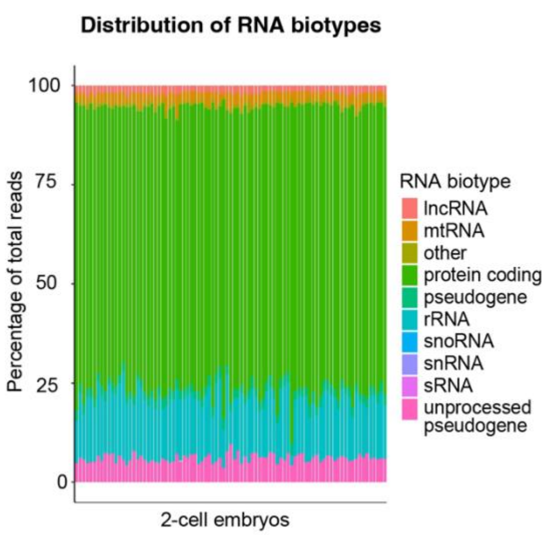


**B**

**TF binding motifs enriched in 2-cell embryos of ELS sEVs-inj. males**

| TF | Motif | P-value | # target seq | % of targets | % of background |
|---|---|---|---|---|---|
| NFY (CCAAT) | 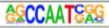 | 1.00E-07 | 11 | 61.1 | 9.3 |
| Klg9 (Zf) | 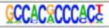 | 1.00E-07 | 11 | 61.1 | 9.3 |
| SP1 (Zf) | 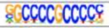 | 1.00E-05 | 11 | 61.1 | 12.4 |
| KLF5 (Zf) | 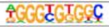 | 1.00E-05 | 14 | 77.8 | 25.4 |
| Sp5 (Zf) | 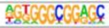 | 1.00E-05 | 14 | 77.8 | 25.4 |
| PRDM15 (Zf) | 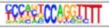 | 1.00E-05 | 9 | 50.0 | 8.5 |
| HIF-1b (HLH) | 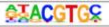 | 1.00E-05 | 11 | 61.1 | 14.7 |
| E2F (E2F) | 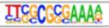 | 1.00E-04 | 5 | 27.8 | 1.8 |
| KLF10 (Zf) | 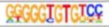 | 1.00E-04 | 8 | 44.4 | 7.7 |
| IRF4 (IRF) | 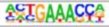 | 1.00E-04 | 5 | 27.8 | 2.4 |
| MyoG (bHLH) | 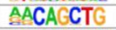 | 1.00E-04 | 8 | 44.4 | 8.6 |
| Hoxa9 | 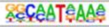 | 1.00E-04 | 10 | 55.6 | 14.8 |
| Sp2 (Zf) | 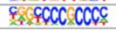 | 1.00E-04 | 14 | 77.8 | 31.6 |
| Ptf1a (bHLH) | 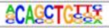 | 1.00E-03 | 11 | 61.1 | 20.8 |
| KLF1 (Zf) | 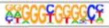 | 1.00E-03 | 11 | 61.1 | 21.0 |
| PRDM1 (Zf) | 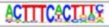 | 1.00E-03 | 5 | 27.8 | 3.4 |
| cMyc (bHLH) | 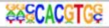 | 1.00E-03 | 5 | 27.8 | 3.7 |
| E2F7 (E2F) | 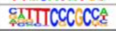 | 1.00E-03 | 5 | 27.8 | 3.8 |

**C**

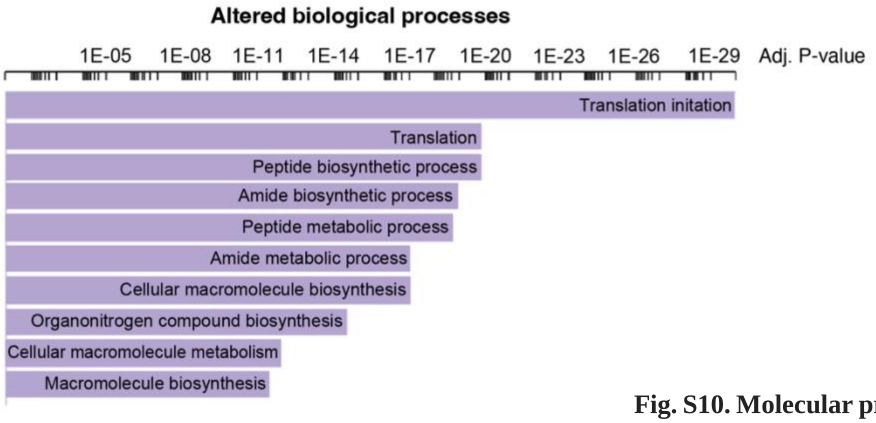


**D**

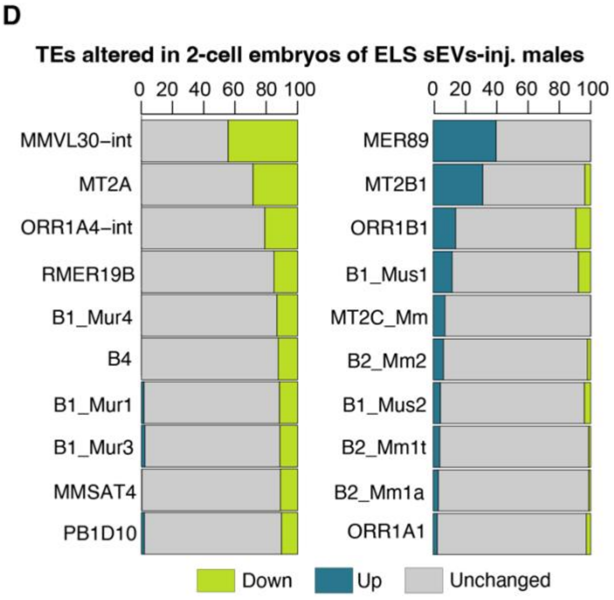


**Fig. S10. Molecular profiles of 2-cell embryos obtained by IVF. A.** Distribution of RNAs biotypes in 2-cell embryos from sperm of control males or males injected with ELS-exposed plasma sEVs at D12 post-injection. *n*=92 embryos. **B.** Transcription factors (TF) and their corresponding binding motifs enriched in differentially upregulated genes in 2-cell embryos from sperm of males injected with ELS-exposed plasma sEVs compared to sperm of males injected with control plasma sEVs. Benjamini-Hochberg test, $P_{adj.}<0.05$. *n*=92 embryos. **C.** Top altered biological processes in genes significantly upregulated in 2-cell embryos from sperm of males injected with ELS-exposed plasma sEVs, compared to controls. Benjamini-Hochberg test, $P_{adj.}<0.0001$. **D.** Down- and up-regulated transposable elements (TEs) in 2-cell embryos identified by RNA-seq. TEs include MMLV30-int 45% of which are significantly down-regulated, and MER89 40% of which are significantly up-regulated in embryos obtained using males injected with plasma sEVs from ELS-exposed mice. Benjamini-Hochberg test, $P_{adj.}<0.05$. *n*=92 embryos.

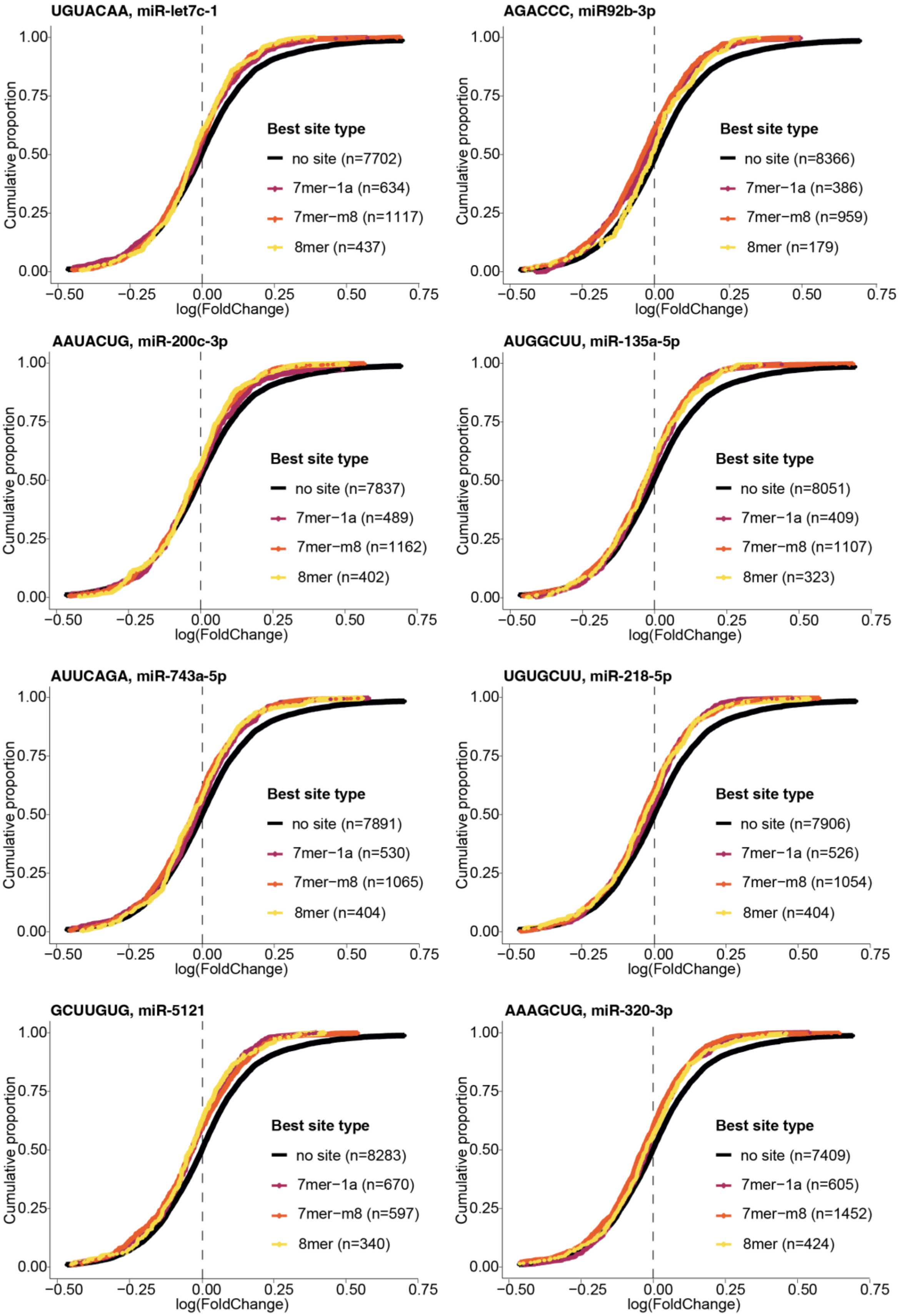


**Fig. S11. Expression of target genes of sperm miRNAs in 2-cell embryos obtained by IVF.** Cumulative distribution plots of all genes' expression in the 2-cell embryos versus expression of genes that have different number of binding sites to miRNAs, that are differentially expressed in the sperm of males injected with ELS-exposed plasma sEVs compared to controls.

**Table S1.**

Full list of RNAs (1.1), lipids (1.2) and metabolites (1.3) altered in sEVs by MSUS.

**Table S2.**

Full metabolic datasets in plasma sEVs isolated by SEC (2.1), plasma from sEVs-injected males (2.2), plasma from D12 offspring (2.3), plasma from D46 offspring (2.4), metabolites that are up- (2.5) or down- (2.6) regulated in plasma from sEVs-injected males.

**Table S3.**

List of small RNAs (3.1) and long RNAs (3.2) altered in sperm at D12, and enriched TF binding sites (3.3). Significant RNAs highlighted in blue.

**Table S4.**

List of small RNAs (4.1) and long RNAs (4.2) altered in sperm at D46. Significant RNAs highlighted in blue.

**Table S5.**

List of RNAs (5.1), enriched components (5.2) and small RNAs (5.3) in embryos derived from sperm from plasma-sEVs injected males.
